# SARS-CoV-2 nucleocapsid protein forms condensates with viral genomic RNA

**DOI:** 10.1101/2020.09.14.295824

**Authors:** Amanda Jack, Luke S. Ferro, Michael J. Trnka, Eddie Wehri, Amrut Nadgir, Xammy Nguyenla, Katelyn Costa, Sarah Stanley, Julia Schaletzky, Ahmet Yildiz

## Abstract

The severe acute respiratory syndrome coronavirus 2 (SARS-CoV-2) infection causes COVID-19, a pandemic that seriously threatens global health. SARS-CoV-2 propagates by packaging its RNA genome into membrane enclosures in host cells. The packaging of the viral genome into the nascent virion is mediated by the nucleocapsid (N) protein, but the underlying mechanism remains unclear. Here, we show that the N protein forms biomolecular condensates with viral genomic RNA both in vitro and in mammalian cells. Phase separation is driven, in part, by hydrophobic and electrostatic interactions. While the N protein forms spherical assemblies with unstructured RNA, it forms asymmetric condensates with viral RNA strands that contain secondary structure elements. Cross-linking mass spectrometry identified a region that forms interactions between N proteins in condensates, and truncation of this region disrupts phase separation. We also identified small molecules that alter the formation of N protein condensates. These results suggest that the N protein may utilize biomolecular condensation to package the SARS-CoV-2 RNA genome into a viral particle.

## Introduction

The SARS-CoV-2 virus consists of a 30 kb single-stranded RNA genome packaged into a 100 nm diameter membrane enveloped virion. SARS-CoV-2 encodes for multiple proteins involved in viral assembly and propagation (*1*) and infects human cells by binding its spike (S) protein to the ACE2 receptor on host cells (*2–4*). While the majority of current efforts to treat COVID-19 have focused on targeting this interaction (*5, 6*), not much work has been done to stop the proliferation of the virus in host cells following infection. Condensation of the viral genome into a virion is primarily driven by the nucleocapsid (N) protein (*7*), which is the most abundant viral protein in infected cells (*8, 9*). A large pool of free N protein is expressed early in infection (*10*) and only a small fraction is transferred into mature virions (*9*). The N protein accumulates at the replication transcription complex (RTC) (*11, 12*) where it enhances replication and the transcription of viral RNA (*13, 14*). The N protein also restructures viral genomic RNA into shell-shaped structures (~15 nm in diameter), which contain approximately 12 N proteins and 800 nucleotides of viral RNA (*7, 15*). These viral ribonucleoprotein complexes (vRNPs) form asymmetric “beads on a string” structures which then bind to the viral membrane (M) protein on the surface of the ER-Golgi intermediate compartment (ERGIC) to trigger the budding of the vRNP complex.

The mechanism by which N remodels the viral RNA and packages it into a viral particle is not well understood. Recent studies proposed that the replication and packaging of viruses involve liquid-liquid phase separation (LLPS) (*16–18*). Biomolecular condensation drives the formation of membrane-less organelles such as the nucleolus, centrosome, stress granules, and P granules through a network of weak and multivalent interactions. Nucleic acids are highly involved in the formation of biomolecular condensates because they can scaffold multivalent interactions (*19–21*). Coronaviruses are involved with phase-separated structures such as stress granules (*10, 22*) and replicate at dynamic clusters associated with the ERGIC, suggesting that phase separation may play a critical role in the replication and packaging of SARS-CoV-2.

Previous studies in HIV-1, negative-sense RNA viruses, and SARS-CoV showed that nucleocapsid proteins drive the formation of phase-separated condensates in the cytosol (*23–26*). The N protein of SARS-CoV-2 also contains many of the characteristic domain features common in phase separating proteins. It contains a well-conserved N-terminal domain (NTD) and a C-terminal domain (CTD, Figure 1A), and 40% of its primary sequence is predicted to be part of intrinsically disordered regions (IDRs, Figure 1B). The NTD (aa: 44-174) interacts nonspecifically with RNA and recognizes a nucleotide sequence in the 3’ end of the viral genome (*13*). The CTD (aa: 257-366) mediates dimerization (*27*), but the N protein can also self-associate into tetramers and higher oligomers (*28, 29*). The NTD and CTD are separated by an IDR that contains a serine/arginine-rich (SR) motif, which has been associated with phase separation in other ribonucleoproteins (*13, 30*). The N and C-terminal IDRs are less conserved but contain arginine- and lysine-rich disordered regions (Figure 1B), which may facilitate additional interactions with the negatively-charged RNA backbone (*31*) and drive biomolecular condensation of RNA (*32, 33*). The N-terminal IDR contains a predicted prion-like domain (PLD, Figure 1A) that can potentially trigger protein demixing (*13, 30, 34*). The C-terminal IDR of SARS-CoV N mediates binding to the M protein (*35, 36*). The N protein is highly positively-charged (+24 in pH 7.4) (*37*), and the CTD and disordered regions also interact with negatively-charged RNA and promote vRNP packaging (*38*). The precise roles of these domains in the phase separation of N protein remain to be elucidated.

**Figure 1.**
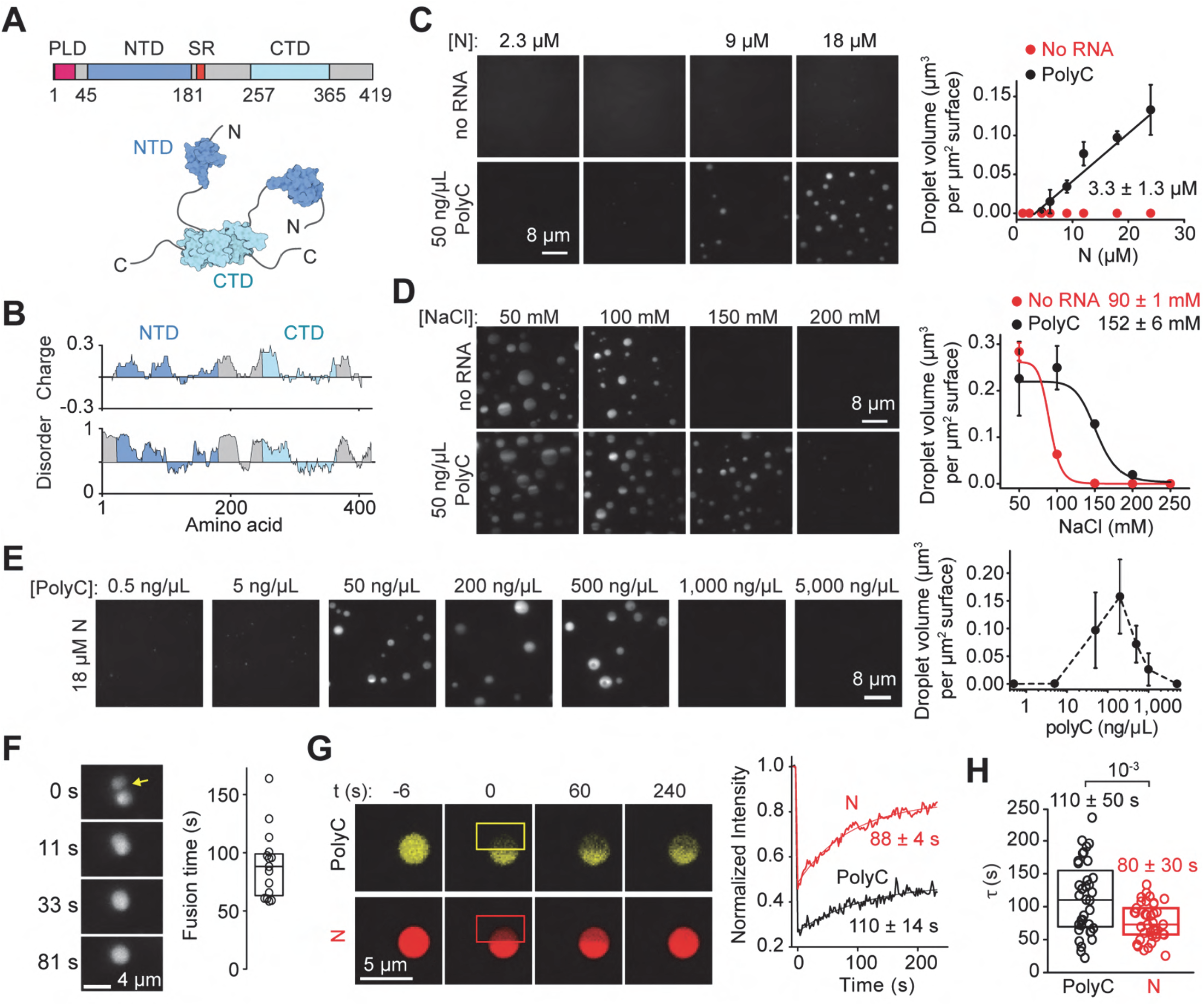
The SARS-CoV-2 N protein phase separates with RNA in vitro. **(A)** Domain organization and the schematic of the N protein dimer. **(B)** Sliding window plot of charge distribution (EMBOSS) and disorder prediction (IUPred2A) for the N protein. Charge y-axis represents mean charge across a 30-residue sliding window. Disorder prediction (1, disordered; 0, ordered) was calculated using the “long disorder” setting, encompassing a 30-residue sliding window. **(C)** (Left) Images of the LD655-labeled N protein in the presence and absence of polyC RNA in 150 mM NaCl. (Right) The total volume of N-RNA condensates settled per micron squared area on the coverslip with 50 ng/μL polyC RNA (mean ± s.d., n = 20 with two technical replicates). A linear fit (solid line) reveals c_sat_ (± s.e.), the minimum N protein concentration for condensate formation (see Methods). **(D)** (Left) Condensates formed by 24 μM LD655-labeled N protein in the presence or absence of 50 ng/μL polyC RNA dissolve by increasing NaCl concentration. (Right) The total volume of N condensates settled per micron squared area on the coverslip with increasing salt concentration (mean ± s.d., n = 20 with two technical replicates). Solid curves represent a fit to a dose-response equation to determine IC_50_ (±s.e.). **(E)** The stoichiometry of the N protein and RNA affects phase separation. (Left) Example pictures show that Cy3-labeled N protein forms condensates with different concentrations of polyC RNA. The N protein concentration was set to 18.5 μM. (Right) The total volume of N-polyC condensates settled per micron squared area on the coverslip under different RNA concentrations (mean ± s.d.; n = 20, two technical replicates). **(F)** (Left) The fusion of N-polyC condensates formed in the presence of 18.5 μM LD655-labeled N and 50 ng/μL polyC RNA. (Right) Fusion time of N-polyC condensates (mean ± s.d., n = 15 fusion events). The center and edges of the box represent the median with the first and third quartiles. **(G)** (Left) Representative FRAP imaging of an N-polyC condensate. The image of a condensate before the time of photobleaching (0 s) shows colocalization of Cy3-polyC and LD655-N in the condensate. Rectangles show the photobleached area. (Right) Fluorescence recovery signals of N and polyC in the bleached region. Solid curves represent a single exponential fit to reveal the recovery lifetime (τ, ±95% confidence interval). **(H)** The distribution of fluorescence recovery lifetimes of N and polyC in droplets (n = 37). The center and edges of the box represent the median with the first and third quartiles. The p-value was calculated from a two-tailed t-test.

In this study, we purified the SARS-CoV-2 N protein from human embryonic kidney (HEK293) cells and observed that N protein forms biomolecular condensates with both homopolymorphic and viral genomic RNA under physiological salt conditions in vitro. The length and structure of the RNA determined the material properties of the N/RNA condensates (*39*). We also showed that the N protein forms liquid condensates in mammalian cells. Cross-linking mass spectrometry (CLMS) identified two regions flanking CTD with interactions enriched within the condensed phase, and the deletion of one of these regions fully abrogated condensate formation in vitro. Together, our results indicate that the N protein phase separates with genomic RNA of SARS-CoV-2, which may play an important role in the packaging of new viral particles in host cells.

## Results

### The N protein phase separates with RNA

We first asked whether the N protein forms bimolecular condensates in the presence or absence of viral RNA in vitro. To address this, we expressed wild-type (WT) N protein in HEK293 cells and purified it in a high salt buffer (1 M NaCl) to eliminate the retention of RNA from human cells (*40*) (Figure 1_figure supplement 1). Consistent with recent studies, purified N protein eluted from gel filtration as an oligomer in 300 mM NaCl (*27*) and had high affinity for binding to various RNA substrates (*41, 42*) (Figure 1_figure supplement 1). The protein was labeled with a fluorescent dye (LD655) at the C-terminal ybbR tag and introduced to a flow chamber in the presence or absence of RNA. Phase separation was monitored by the settling of N or N-RNA condensates on to the coverslip within 25 minutes under highly inclined and laminated optical sheet (HiLO) excitation (see Methods for details). In the absence of RNA, N protein did not form condensates in physiological salt (150 mM NaCl) (Figure 1C). Similarly, 2-kb long polyC RNA homopolymer did not form any condensates in the absence of N protein (Figure 1_figure supplement 2A-B). Mixing of 50 ng/μL polyC RNA and the N protein resulted in the formation of condensates in 150 mM NaCl (Figure 1C). LD655-N and Cy3-polyC colocalized well in the droplets, suggesting that phase separation of N protein is driven by RNA (Figure 1_figure supplement 2C). The analysis of the condensates settled on the coverslip revealed a saturation concentration (c_sat_) of 3.3 ± 1.3 μM for N protein in the presence of 50 ng/μL polyC RNA (± s.e., Figure 1C), consistent with the abundance of N protein in infected cells (*43*). The partition coefficient of N protein into condensates was 13 ± 2 (±s.d.).

Although N protein is unable to phase separate without RNA in 150 mM salt, 24 μM N protein efficiently formed condensates at lower salt with a half-maximal inhibition constant (IC_50_) of 90 ± 1 μM NaCl (±s.e.) (Figure 1D). The addition of 50 ng/μL polyC RNA increased IC_50_ to 152 ± 6 mM NaCl (Figure 1D). The ability of N protein to phase separate without RNA and sensitivity of these condensates to salt shows that they are dynamic structures driven, in part, by electrostatic and pi interactions among N proteins, as observed for other proteins that phase separate (*44, 45*).

Condensate formation of N protein and RNA was dependent on protein-RNA stoichiometry. At 18 μM N protein, condensate formation was not observed in the presence of 0-5 ng/μL polyC RNA. Increasing the RNA concentration promoted phase separation with an optimal RNA concentration of 100-500 ng/μL, in which charge neutralization occurs with the positively-charged N protein (*46*). A further increase in RNA concentration dissolved these condensates (Figure 1E) (*47, 48*). This reentrant phase separation behavior is characteristic of heterotypic RNA and protein interactions in phase separating systems (*47*).

We also showed that N-polyC condensates exhibit liquid-like, rather than solid-like, material properties. First, these condensates were nearly spherical with an aspect ratio of 1.3 ± 0.6 (mean ± s.d.). Second, we observed the fusion of condensates with a mean fusion time of 90 ± 30 s (mean ± s.d.) after they come into contact (Figure 1F). Finally, fluorescence recovery after photobleaching (FRAP) experiments showed that 90 ± 10% of N protein in condensates can slowly exchange with the solvent with the half recovery time of 80 ± 30 s (mean ± s.d.). In comparison, polyC RNA exhibited slower recovery and a lower mobile fraction (Figure 1G-H, Figure 1_figure supplement 3), indicating that the RNA may be stabilized by a network of interactions with multiple N proteins in the condensates.

Similar to polyC RNA, polyA and polyU RNA substrates produced spherical condensates at micromolar concentrations of the N protein (Figure 2A). However, combining N protein with polyG that forms G-quadruplexes or polyAU that forms base-pair interactions led to the formation of non-spherical condensates (Figure 2A). These results indicate that the N protein forms liquid condensates with nonspecific RNA that do not contain RNA sequence-specific secondary structures, whereas RNA substrates that form internal base-pairing interactions result in asymmetric condensates (*19, 39, 49, 50*).

**Figure 2.**
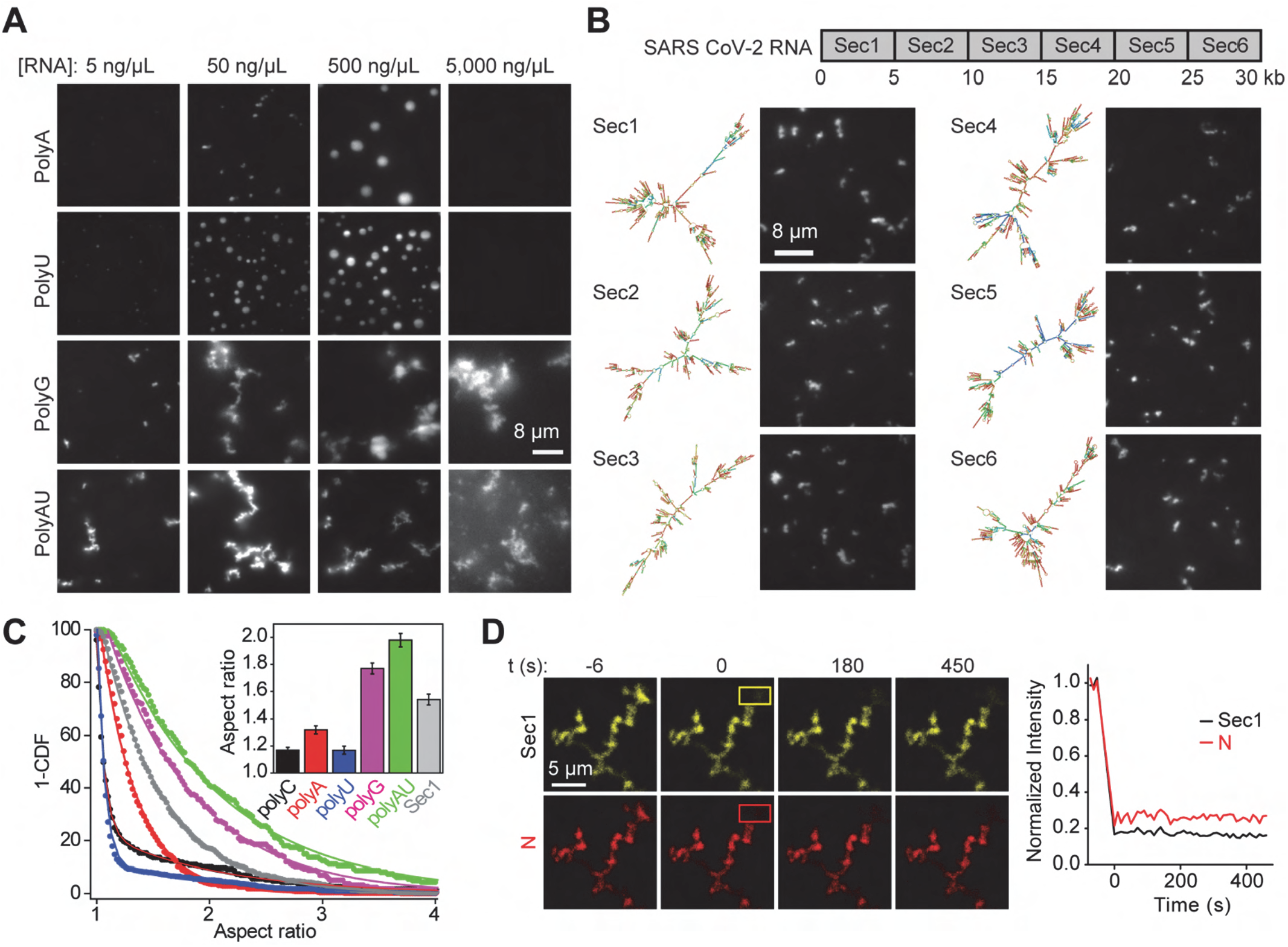
The N protein forms asymmetric condensates with viral RNA. **(A)** The N protein forms spherical condensates with unstructured RNA (polyA and polyU) but forms asymmetric condensates with structured RNA (polyG and polyAU). The N protein concentration was set to 18.5 μM. **(B)** (Top) SARS-Cov-2 genomic RNA was divided into 6 sections. (Bottom, left) Structure prediction of each section and (Bottom, right) the formation of asymmetric N condensates in the presence of 18 nM RNA. The N protein concentration was set to 18.5 μM. **(C)** The inverse cumulative distribution (1-CDF) of the aspect ratio of individual N condensates formed with different RNA substrates. The concentrations of N protein, RNA homopolymers, and Sec1 RNA were set to 18.5 μM, 50 ng/μL, and 18 nM, respectively. Solid fits represent a fit to exponential decay. (Insert) Decay constants of the exponential fits (±s.e.). **(D)** (Left) Representative FRAP imaging of an N-Sec1 condensate. The image of a condensate before the time of photobleaching (0 s) shows colocalization of Cy3-Sec1 and LD655-N in the condensate. Rectangles show the photobleached area. (Right) N and Sec1 do not exhibit fluorescence recovery in the bleached region (n = 16).

Next, we sought to characterize how the N protein interacts with SARS-CoV-2 genomic RNA. The SARS-CoV-2 RNA genome was reverse transcribed and assembled into a DNA plasmid (*51*). Using this plasmid, we generated six 5 kb and two 1 kb fragments of the viral RNA genome via in vitro transcription (*52*). Unlike polyC, in silico methods predict that these RNA fragments can form intra- and inter-molecular base-pairing interactions and contain extensive secondary structure elements (Figure 2B, Figure 2_figure supplement 1A) (*39, 53, 54*). Similar to synthetic RNA substrates that form base-pair interactions (Figure 2A), the N protein formed non-spherical condensates with viral RNA fragments in vitro (Figure 2B, Figure 2_figure supplement 1A-B). The aspect ratio of condensates formed with viral RNA was higher than unstructured polymorphic RNA but lower than those formed with polyG and polyAU fragments (Figure 2C, Figure 2_figure supplement 1C). This could be due to the presence of both structured and unstructured segments within the viral RNA (Figure 2B). These asymmetric structures formed across a wide range of protein and viral RNA concentrations and were dissolved by increasing the salt concentration, but they did not change shape over time or fuse with each other and were not strongly affected by raising the temperature from 20 °C to 37 °C (Figure 2_figure supplement 2). We also did not detect fluorescence recovery of either the LD655-labeled N protein or Cy3-labeled viral RNA in FRAP assays (Figure 2D, Figure 2_figure supplement 3). We concluded that N protein forms solid-like condensates with viral RNA in vitro and that intramolecular interaction of the RNA influences the material properties of the condensate (*19, 39, 49*).

### Cross-linking mass spectrometry identifies N protein interaction sites

To understand the mechanism of phase separation of the N protein, we performed CLMS to identify interactions between different domains of the full-length N protein in the absence of RNA (*55*). CLMS detects protein-protein contacts by covalently capturing nearby residues with bifunctional reagents. We first crosslinked the soluble (not phase separated) N protein in 300 mM KAc using a bifunctional crosslinker bis(sulfosuccinimidyl) suberate (BS3) (Figure 3_figure supplement 1A). We detected that the N-terminal half of the protein, including NTD, makes diverse contacts throughout the entire protein (Figure 3_figure supplement 1A). There was also an abundance of contacts between the regions immediately flanking CTD on either side, referred to as R1 (aa: 235-256) and R2 (aa: 369-390).

Next, we performed quantitative CLMS measurements (*56*) comparing the soluble N protein in 300 mM KAc with phase-separated N protein in 100 mM KAc (Figure 3A). The soluble N protein was crosslinked with heavy (D12) BS3 whereas the phase-separated protein was crosslinked with light (H12) BS3. As a result, the crosslinked precursor ions from high and low salt conditions were spaced by 12 Da (Figure 3A). Interactions between specific regions that promote condensate formation are implicated by the ratio of the crosslinked precursor ion signal and its corresponding isotopic doublet (Figure 3A) (*56*). This experiment was repeated by reversing the labels, such that (H12) BS3 was used to crosslink the soluble N protein and (D12) BS3 was used for the phase-separated N protein. Across two independent experiments, 29 unique crosslinks were enriched and 30 crosslinks were depleted upon phase separation (Figure 3B, Figure 3_figure supplement 1B). Remarkably, the analysis of the crosslink fold change (Table S1) revealed that nearly all of the enriched interactions are concentrated in regions R1 and R2, whereas depleted crosslinks more uniformly spanned the entire primary sequence (Figure 3C). These results suggest that the interactions involving the amino acids in regions R1 and R2 may be important in driving phase separation (Figure 3D, Figure 3_figure supplement 1C).

**Figure 3.**
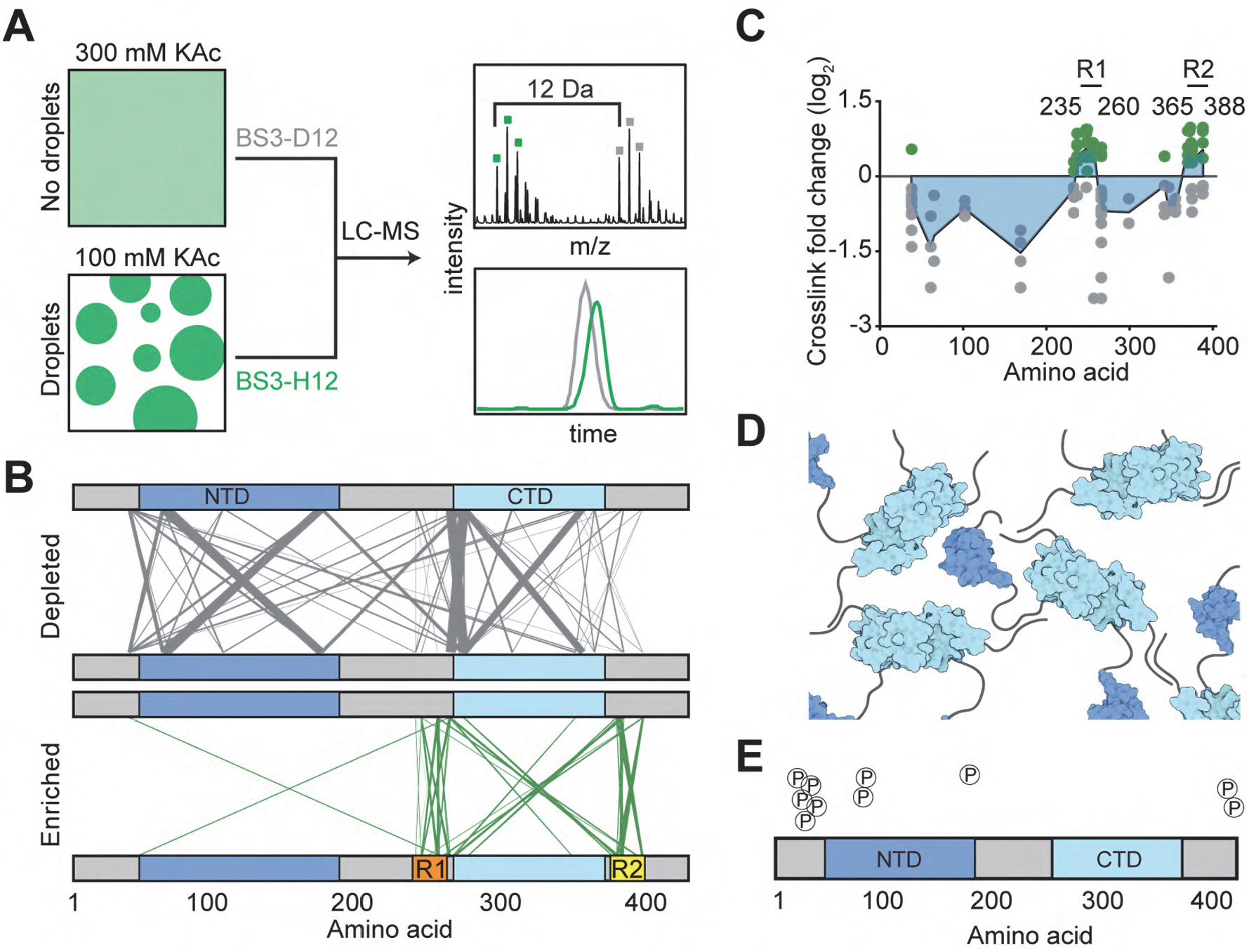
CLMS reveals inter-domain interactions of the N protein. **(A)** Schematic of the CLMS experiment. (Left) A high salt (300 mM KAc) buffer disrupts N condensates, whereas a low salt (100 mM KAc) buffer promotes condensate formation. (Right, top) Example of an individual crosslinked peptide in quantitative CLMS analysis. Precursor ions from the high salt (gray) and low salt (green) BS3 crosslinking conditions show the 12 Da shift between light (H12) and heavy (D12) crosslinkers. (Right, bottom) Ion chromatograms from the first three isotopes of each doublet were extracted and expressed as the ratio of peak areas. **(B)** The plot of crosslinks depleted and enriched in the condensate condition. The width and transparency of the lines scale with the number of times the crosslink was detected across 3 independent experiments. **(C)** Fold changes of crosslink abundance upon condensate formation of N. As crosslinks contain two positions, fold change information is plotted at both positions. Only crosslinks with p-values less than 0.05 are included. Green and grey dots represent crosslinks enriched and depleted in the condensate condition, respectively. The blue area represents a plot of median crosslink fold change. **(D)** Model for how multiple N dimers could phase separate via their disordered regions. **(E)** Phosphorylation sites detected by the CLMS experiment in 300 mM KAc.

Our mass spectrometry (MS) analysis also found phosphorylation sites on the N protein (Figure 3E). While some of these sites have been identified in previous studies (*10, 57, 58*), we also identified several novel sites (Table S2). Although one of the phosphorylation sites (S176) is involved in a crosslink, the phosphorylated and unphosphorylated peptides were both strongly depleted in condensates, suggesting that S176 phosphorylation does not play a major role in phase separation (Tables S1 and S2, Figure 3_figure supplement 1B). Additionally, MS identified native proteins that co-purified with the N protein in 1 M salt (Table S3). Consistent with the recruitment of N to stress granules in cells (*13, 46, 59*), two of the most frequently identified proteins were stress granule proteins G3BP1 and G3BP2, with R2 of the N protein interacting with G3BP1 (Figure 3_figure supplement 1D).

### The C-terminal region and phosphorylation modulate phase separation

The quantitative CLMS experiments show that interactions between R1 and R2 are correlated with the formation of condensates. However, we could not exclude the possibility that some of the changes in pairwise interactions are due to differences in protein structure or electrostatic interactions induced by differences in salt concentrations used for the soluble and condensate phase. To directly test the predictions of the CLMS results, we determined how different domain deletions affected phase separation of the N protein with polyC RNA under the same salt concentration (Figure 4A, Figure 4_figure supplement 1). Phase separation was only moderately reduced in the ΔSR, ΔPLD, and ΔR1 mutants (Figure 4B, Figure 4_figure supplement 2), suggesting that these regions are not essential for phase separation. In comparison, deletion of the R2 sequence fully abolished the formation of condensates with polyC RNA (Figure 4B, Figure 4_figure supplement 2A-D). Similarly, ΔR2 was unable to form condensates with viral RNA, whereas other deletion mutants phase separated with the same RNA fragments (Figure 4_figure supplement 2E). ΔR2 maintained a high affinity to bind polyC and viral RNA (Figure 4_figure supplement 1C), excluding the possibility that the disruption of phase separation is due to the lack of protein-RNA interactions. These results show that R2 primarily drives phase separation of N protein with RNA, consistent with the proposed role of this sequence in the oligomerization of N protein (*27*).

**Figure 4.**
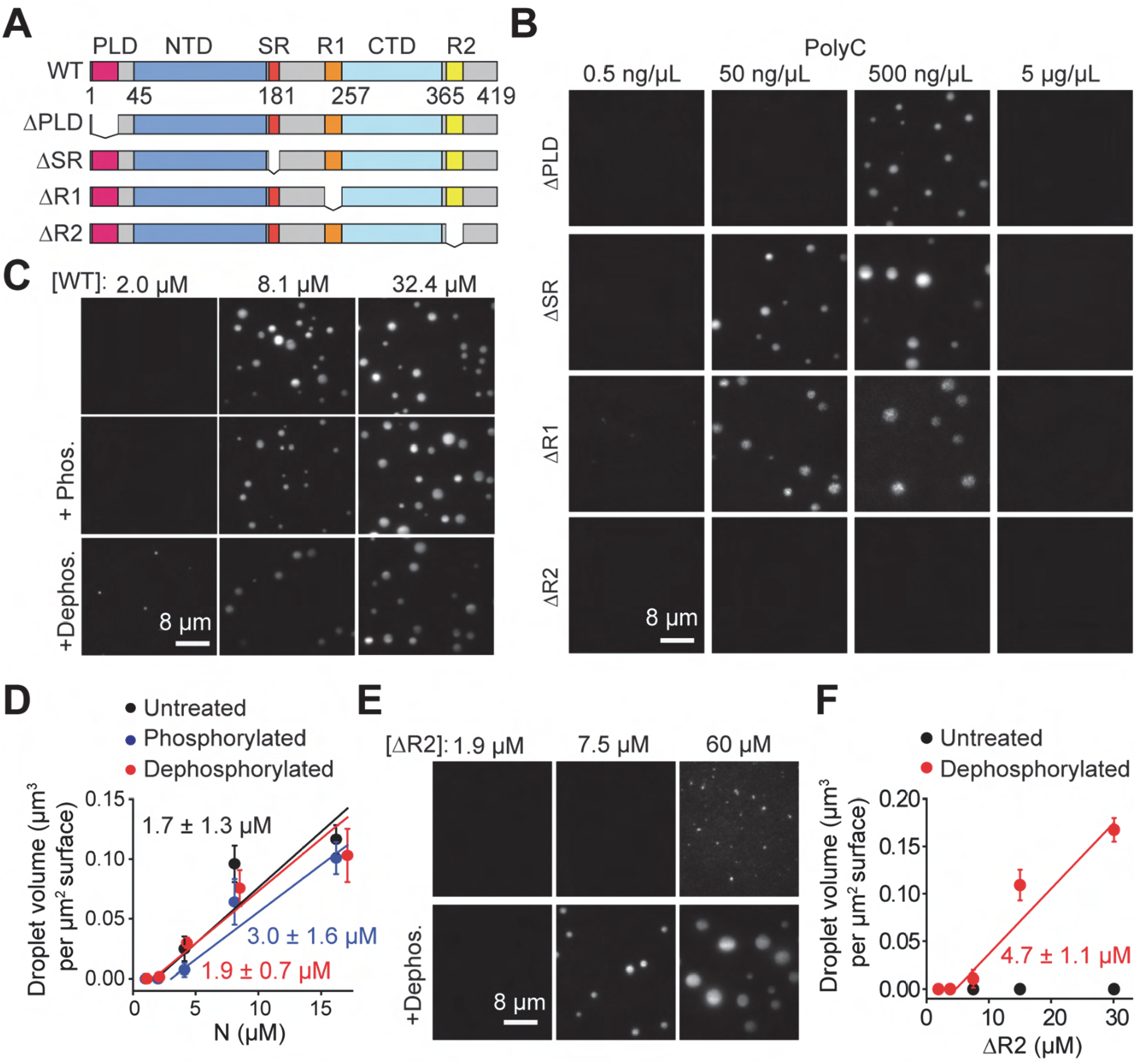
The effect of domain truncations and phosphorylation on phase separation of the N protein. **(A)** Truncation mutants of the N protein. **(B)** While ΔPLD, ΔSR, and ΔR1 phase separate, ΔR2 does not phase separate when mixed with polyC RNA. Protein concentration was set at 18 μM for all conditions. **(C)** Images of condensates formed by untreated, phosphorylated, and dephosphorylated N protein in 50 ng/μL polyC RNA. **(D)** The total volume of N-RNA condensates settled per micron squared area on the coverslip under different phosphorylation conditions (mean ± s.d., n = 20 with two technical replicates). Linear fits (solid lines) reveal c_sat_ (± s.e.). **(E)** Images of condensates formed by untreated and dephosphorylated ΔR2 in 50 ng/μL polyC RNA. **(F)** The total volume of ΔR2-RNA condensates settled per micron squared area on the coverslip as a function of ΔR2 concentration (mean ± s.d., n = 20 with two technical replicates). The linear fit (solid line) reveals c_sat_ (± s.e.).

Recent studies proposed that the N protein is phosphorylated early in infection, and this results in localization of N protein with the RTC, where it enhances transcription of subgenomic RNA (*60*). However, nucleocapsid formation does not require phosphorylation and N protein in SARS-CoV viruses are hypophosphorylated (*60, 61*). The underlying molecular mechanism and kinases and phosphatases responsible for post-translational modification of N protein remain poorly understood (*8*). To understand how phosphorylation affects phase separation, we treated the full-length N protein with casein kinase 2 and λ protein phosphatase (see Methods). While kinase treatment did not alter the migration of N protein on a denaturing gel, phosphatase treatment resulted in a reduction in molecular weight (Figure 4_figure supplement 1A), suggesting that the N protein expressed in human cells is phosphorylated. Phosphatase treatment resulted in phase separation at slightly lower concentrations in comparison to kinase-treated N protein (Figure 4C-D). Similar to WT, dephosphorylated ΔSR or ΔR1 had only a moderate increase in phase separation (Figure 4_figure supplement 1 and 3). Unlike untreated ΔR2, dephosphorylated ΔR2 exhibited robust phase separation with polyC RNA (Figure 4E,F). These results suggest that phosphorylation negatively regulates phase separation of the N protein, but interactions formed by R2 supersede this inhibition.

### Targeting phase separation of the N protein with small molecules

We next sought to identify small molecules that could interfere with the phase separation of the N protein. The condensates formed by the N protein in the absence or presence of polyC or viral RNA dissolve with the addition of 10% 1,6 hexanediol, indicating that phase separation is driven, at least partially, by hydrophobic interactions (22) (Figure 5A). In comparison, lipoic acid that increases the liquidity of stress granules (*62*) did not affect phase separation (Figure 5A). Using a high-throughput microscopy platform, we also tested whether any of the 1200 compounds in the FDA-approved library alters phase separation of the N protein with polyC RNA in vitro. While none of the compounds fully dissolved the condensates, we identified several compounds that affected their number, size, and shape (Table S4). Nelfinavir mesylate and LDK378 produced larger but fewer condensates, whereas crystal violet, tolcapone, and chlorhexidine enhanced phase separation by increasing the number and size of the condensates (Figure 5B, Figure 5_figure supplement 1A-B). Nilotinib resulted in a 50% increase in condensate volume and altered the shape of the condensates (Figure 5B), which may be due to changes in condensate fusion or maturation. While most drugs did not have a substantial effect on condensates formed by N and viral genomic RNA in vitro, nilotinib addition resulted in the formation of thread-like filaments (Figure 5_figure supplement 1C-D). Morphologically, these filaments were similar to those formed during the liquid-to-solid transition of FUS condensates (*63*), suggesting that nilotinib increases the viscosity of the condensates.

**Figure 5.**
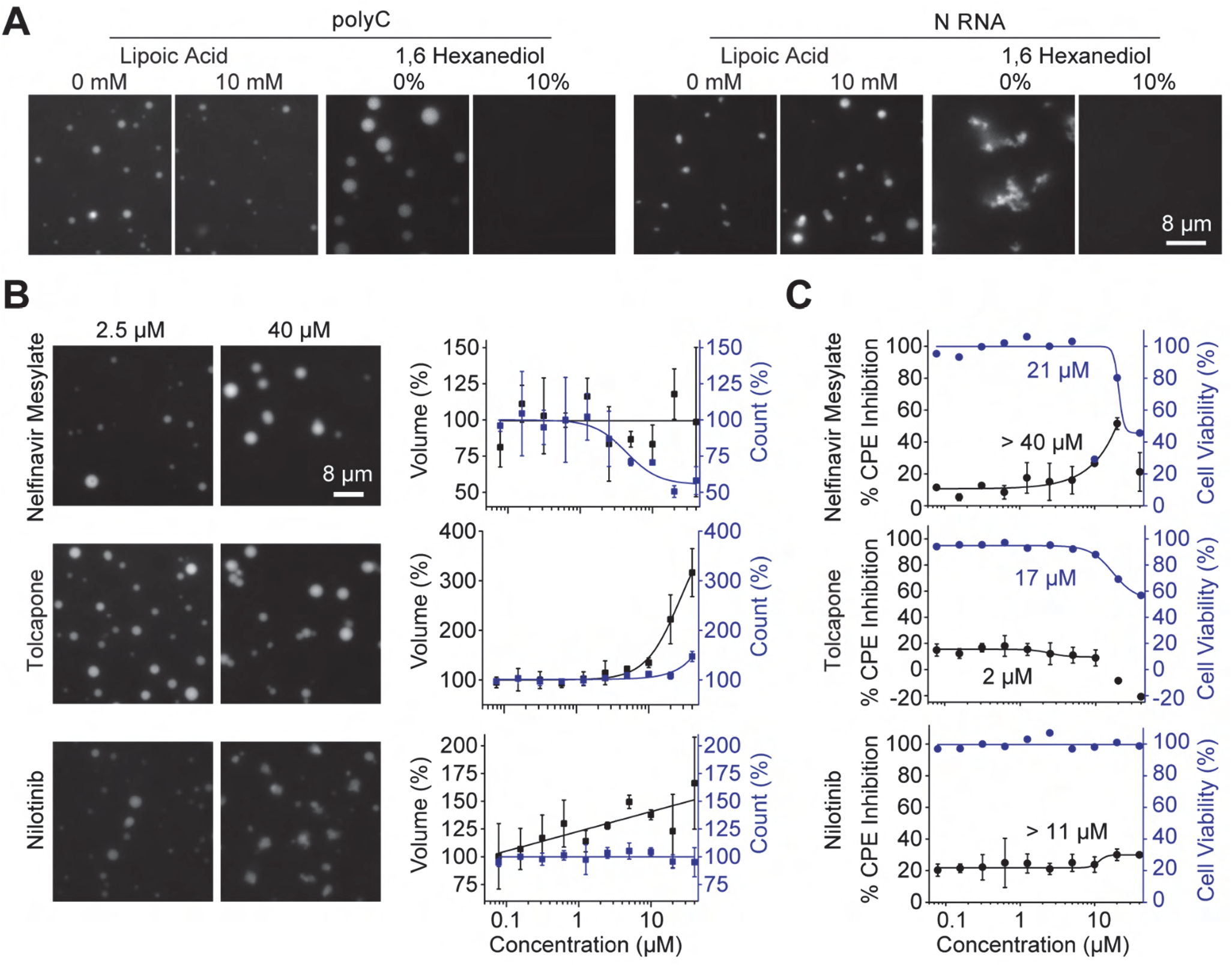
Identification of small molecules that alter phase separation of N in vitro and in vivo. **(A)** Condensates formed in the presence of 7.8 μM N and either 50 ng/μL polyC or 18 nM Sec1 RNA were not affected by 10 mM lipoic acid, but dissolved in the presence of 10% 1,6 hexanediol. **(B)** (Left) Examples of Class I, II, and III drugs. The N protein and polyC RNA concentrations were set to 7.8 μM and 50 ng/μL, respectively. (Right) The percent change on the number (blue) and total volume (black) of N-polyC condensates settled per micron squared area on the coverslip under different drug concentrations (mean ± s.d., n = 8 with two technical replicates). Solid curves represent a fit to a dose-response equation (see Methods). **(C)** Percent CPE inhibition (black, mean ± s.d., two technical replicates) and cell viability (blue, mean) of SARS-CoV-2 infected Vero-E6 cells treated with serial dilutions of drugs. Solid curves represent a fit to a dose-response equation (see Methods) to determine a half-maximal response constant EC_50_ (Table S4).

We also tested these drugs in the human pulmonary epithelial (Calu-3) and the African Green Monkey kidney (Vero-E6) cells infected with SARS-CoV-2 (see Methods), as these cell lines supported high levels of infection (*64*). The cells were treated with different concentrations of drugs 1 hour before infection and the inhibition of SARS-CoV-2-mediated cell death under drug treatment was quantified using the cytopathic effect (CPE) inhibition assay (*64*). The toxicity of the drugs was quantified by measuring the viability of uninfected cells under drug treatment. Among the molecules we identified, nelfinavir mesylate resulted in the highest percent CPE inhibition in both cell lines without significantly affecting cell viability (Figure 5C, Figure 5_figure supplement 2), with the exception that the highest dose (40 μM) was toxic to the cells. Collectively, our drug screen identified compounds that affect the condensate formation of N protein and RNA *in vitro* and inhibit virus-mediated cell death.

### The N protein phase separates in mammalian cells

To test whether N protein also phase separates in mammalian cells, we expressed WT-GFP in HEK293T cells. We observed the formation of distinct puncta in the cytoplasm in WT-GFP expressing cells, which were not observed in control cells expressing GFP only (Figure 6A). FRAP imaging of these puncta revealed a recovery signal with 60% mobile fraction and 6.3 ± 0.1 s recovery lifetime, suggesting that N protein is capable of forming liquid condensates in cells (Figure 6B, Figure 6_figure supplement 1). The recovery of WT-GFP was substantially slower than GFP only (Figure 6C-E), but an order of magnitude faster than that of N-polyC condensates in vitro (Figure 1H) (*8, 40*), suggesting that phase separation of N protein may be regulated inside cells. This difference is not due to the absence or low stoichiometry of RNA in N-GFP puncta in live cells, because N condensates formed in the absence of RNA in vitro also exhibited an order of magnitude slower recovery than N-GFP puncta in live cells (Figure 6_figure supplement 2). We next tested whether deletion of the R2 region disrupts phase separation of N protein in cells. Surprisingly, cells expressing the R2 deletion mutant (ΔR2-GFP) still formed puncta and did not display significantly different FRAP dynamics than WT-GFP in live cells (Figure 6). This may be due to macromolecular crowding of the cellular environment because the addition of 10% PEG triggered phase separation of ΔR2 without RNA in vitro (Figure 6_figure supplement 3). Collectively, these results show the N protein is capable of liquid-liquid phase separation in cells. Differences between in vitro and in vivo phase separation might be attributed to macromolecular crowding of the cytosol or the interaction of the N protein with other cellular proteins, such as G3BP1 (Figure 3_figure supplement 1D) (*46, 59, 65*).

**Figure 6.**
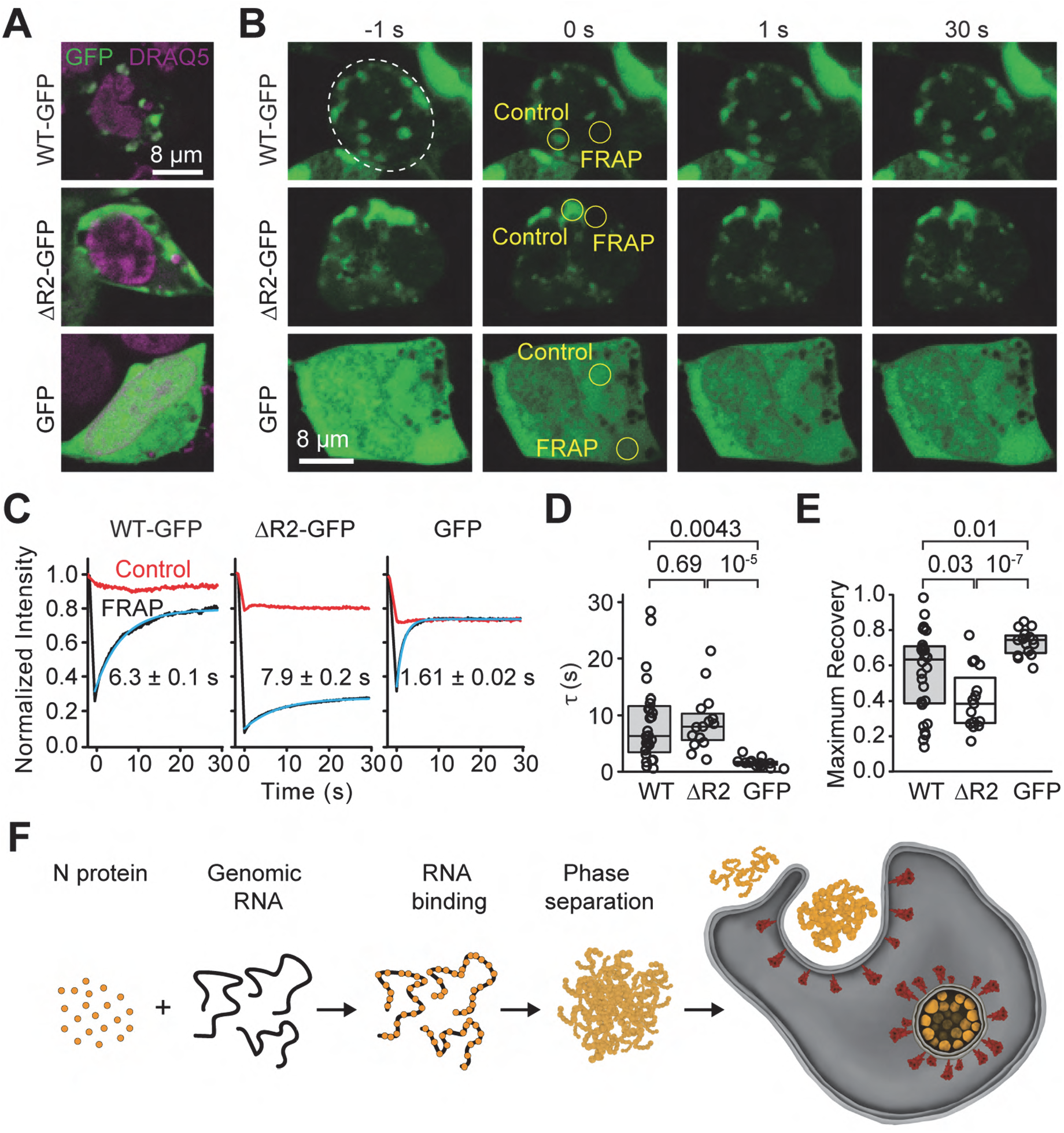
Dynamics of N condensates in vivo. **(A)** Example images of cells expressing N-GFP, ΔR2-GFP, and GFP stained with DRAQ5. **(B)** Representative FRAP imaging of cells exhibiting N-GFP or ΔR2-GFP puncta or high GFP expression. Circles show the photobleached area. **(C)** Fluorescence recovery signals of the protein in the bleached versus the control regions. The solid curve represents a single exponential fit to reveal the recovery lifetime (τ, ±95% confidence interval). **(D)** The distribution of fluorescence recovery lifetimes of cells expressing WT-GFP (n = 28) or ΔR2-GFP (n = 15) exhibiting puncta and cells with high expression of GFP only (n = 15). The center and edges of the box represent the median with the first and third quartiles. The p values were calculated from a two-tailed t-test. **(E)** The maximum fractional recovery after photobleaching of cells expressing WT-GFP (n = 28), ΔR2-GFP (n = 15) or GFP-only (n = 15). The center and edges of the box represent the median with the first and third quartiles. The p values were calculated from a two-tailed t-test. **(F)** Model for remodeling of viral RNA genome by the SARS-CoV-2 N protein. The N protein packages viral genomic RNA through phase separation, which may facilitate efficient replication of the genomic RNA and formation of the enveloped virus.

## Discussion

In this study, we showed that the SARS-CoV-2 N protein phase separates with nonspecific RNA sequences in vitro and the viscosity and shape of these condensates depend on the structure of the RNA substrate. Concurrent studies have shown that the SARS-CoV-2 N protein expressed in bacteria forms biomolecular condensates with RNA in vitro (*8, 32, 40, 46, 50, 52, 66*). We purified N protein from mammalian cells to recapitulate the post-translational modifications that occur in infected human cells (*10*). Consistent with Carlson et al. and Chen et al., we observed that the N protein requires unstructured RNA to form liquid condensates in physiological salt (*8, 66*). Condensates of N with unstructured RNA recovered from photobleaching and relaxed to a spherical shape, suggesting that they are dynamic, liquid-like compartments. In agreement with Iserman et al., we found that RNA structure affects the material properties of the condensate (*52*); base-paired or G-quadruplex-containing RNA produces irregularly-shaped condensates (*50*). N protein also formed solid-like condensates with long fragments of SARS-CoV-2 genomic RNA. These condensates had asymmetric shapes and did not relax over time or with increased temperature (*8, 46*). Viral RNA may drive the formation of abnormal condensate shapes by forming networks of intermolecular interactions, as previously reported (*19, 39, 49, 67*).

We also investigated the mechanisms underlying condensate formation using CLMS and protein engineering. Previous studies reported that prion-like and SR motifs are common features of phase separating proteins (*30, 34*), and the linker sequence outside the SR motif (amino acids 210-247) is essential for phase separation of N expressed in bacteria (*40*). The SR motif was required for N/N interactions in SARS-CoV and for forming puncta in SARS-CoV infected cells (*68*). In addition, the SR motif is highly phosphorylated in human cells, and phosphorylation has been shown to promote interactions with host proteins, nuclear targeting, and transcription of the viral genome, and either enhances or inhibits oligomerization in SARS-CoV (*13, 60, 61, 69*). In SARS-CoV-2, SR phosphorylation is reported to make N condensates more liquid (*8*). However, we observed that the SR region is not necessary for the condensate formation of full-length N protein expressed in human cells. Although the deletion of the entire C-terminal IDR enhances phase separation (*8, 27*), we found that deletion of the R2 motif in this region fully disrupts phase separation of N protein with both polymorphic and viral RNA in vitro. The C-terminal region interacts with the M protein, which was shown to drive phase separation of the N protein in the absence of RNA (*27*). Because the C-terminal region plays an important role in phase separation, binding of the N protein to the membrane-associated M protein through this region can alter the material properties of the N-RNA condensates and initiate virion assembly.

It remains to be demonstrated whether condensate formation of N protein and the viral genome is essential for the propagation of SARS-CoV-2 in human cells. Phase separation has been implicated as a possible mechanism in the viral life cycle. For example, several viruses have been shown to replicate in viral inclusion bodies that are characterized as phase-separated condensates (*24, 25*). A recent in vitro work has shown that nucleoproteins and phosphoproteins of the measles virus form liquid-like membraneless organelles and triggered nucleocapsid formation (*18*), suggesting that phase separation could be a general mechanism for viral replication. In the case of SARS-CoV-2, phase separation of N protein can form membrane-less compartments and function as a selectivity barrier to control the entry of certain agents into these compartments to enhance the efficiency of replication, while sequestering the viral assemblies from the immune response of the host cell (*8*). For example, N protein interacts with stress granule proteins (*50, 59*) and inhibits stress granule formation in uninfected cells (*65, 70*). It remains to be determined whether N protein suppresses stress granule formation in infected cells and whether this plays a role in the efficiency of viral replication. Similarly, recent in vitro studies have shown that N-RNA condensates recruit the components of the SARS-CoV-2 replication machinery (*46*). This mechanism may increase the efficiency of replication of the viral genomic RNA by increasing the local concentration of the replication machinery at the RTC complex.

The phase separation of N protein with viral RNA may also facilitate the compaction and packaging of the genome into the nascent viral particle (Figure 6F). Consistent with this possibility, recent studies have shown that phase separation plays an essential role in the formation of heterochromatin regions in the nucleus of mammalian cells (*71, 72*). However, this model raises several challenges. Because N protein can phase separate with nonspecific RNA substrates, it remains unclear how these condensates may exclude subgenomic viral RNA and other RNA from the host cell. Recent in vitro work has shown that N protein has a higher affinity to bind 5’ and 3’ untranslated regions of the genomic RNA (*52*), which may serve as a mechanism to exclude other RNA from the condensates. In addition, condensates observed in vitro and in vivo are large structures that can potentially contain thousands of N proteins and RNAs, and it is not clear how a virion with a single genomic RNA can bud from these structures. A recent modeling study proposed that the presence of high-affinity sites in genomic RNA can trigger the formation of single-genome condensates (*32*). Alternatively, interaction with the M protein or dephosphorylation of N protein, which has been proposed to trigger the liquid-to-solid transition of N-RNA condensates, at the viral assembly sites may trigger budding of a single genomic copy in virions. Testing of these models requires in-depth studies of N protein and genomic RNA in SARS-CoV-2 infected cells.

Phase separation could also provide a macroscopic readout to study N protein and RNA interactions (*62*) and suggest novel strategies to disrupt genome packaging and viral propagation in infected cells. We performed a screen of an FDA-approved library and identified several compounds that altered the size, number, and shape of N/RNA condensates in vitro. In particular, nelfinavir mesylate binds the SARS-CoV-2 protease (*73–76*) and blocks SARS-CoV viral production (*77*). We showed that nelfinavir mesylate inhibits the proliferation of the virus in the host cell, which may be related to changes it induces on N condensates. Future work in infected cells is needed to address whether nelfinavir mesylate reduces virus-mediated cell death by interfering with the functions of N-RNA condensates.

---

**Table S1.**
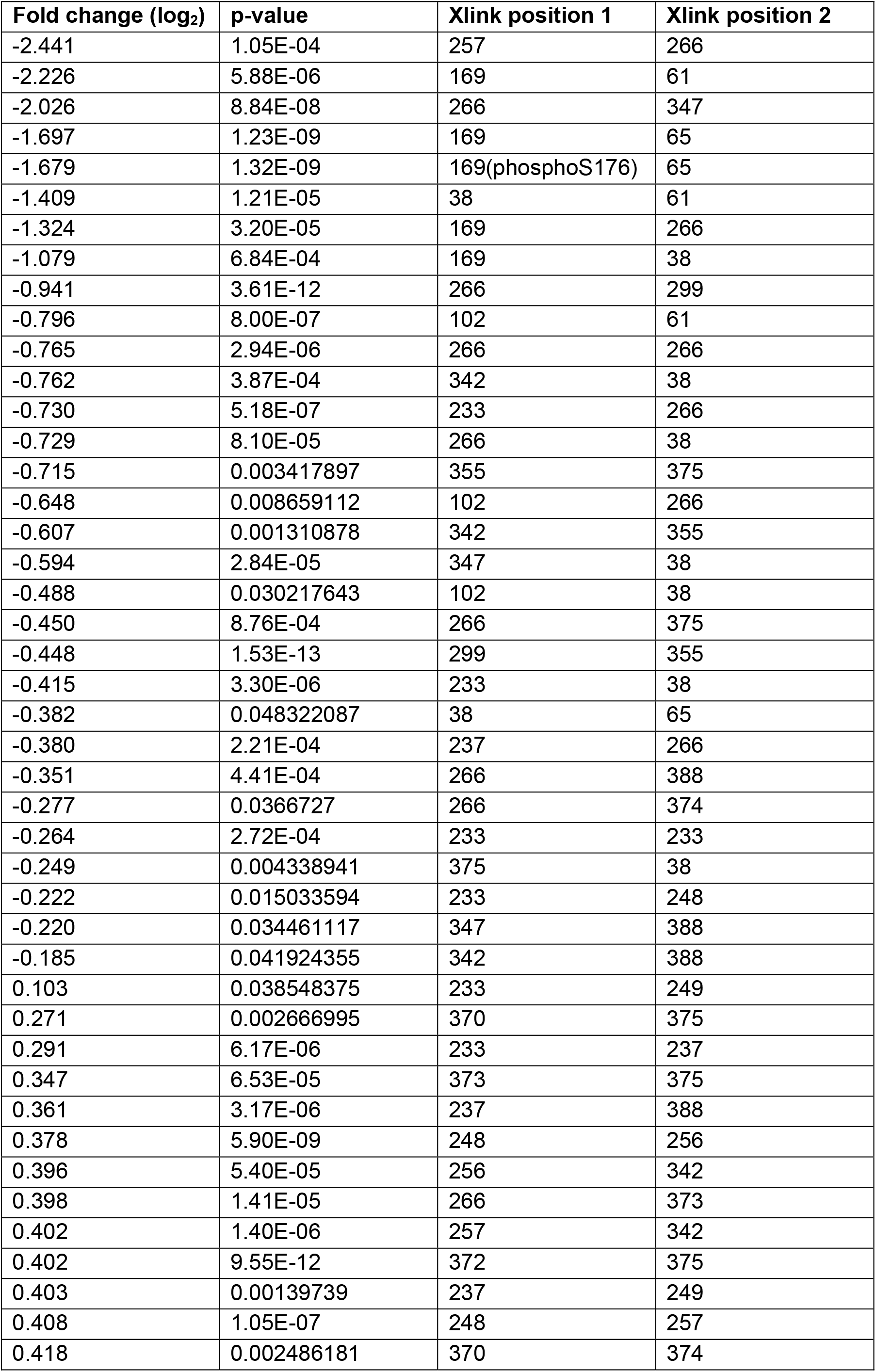
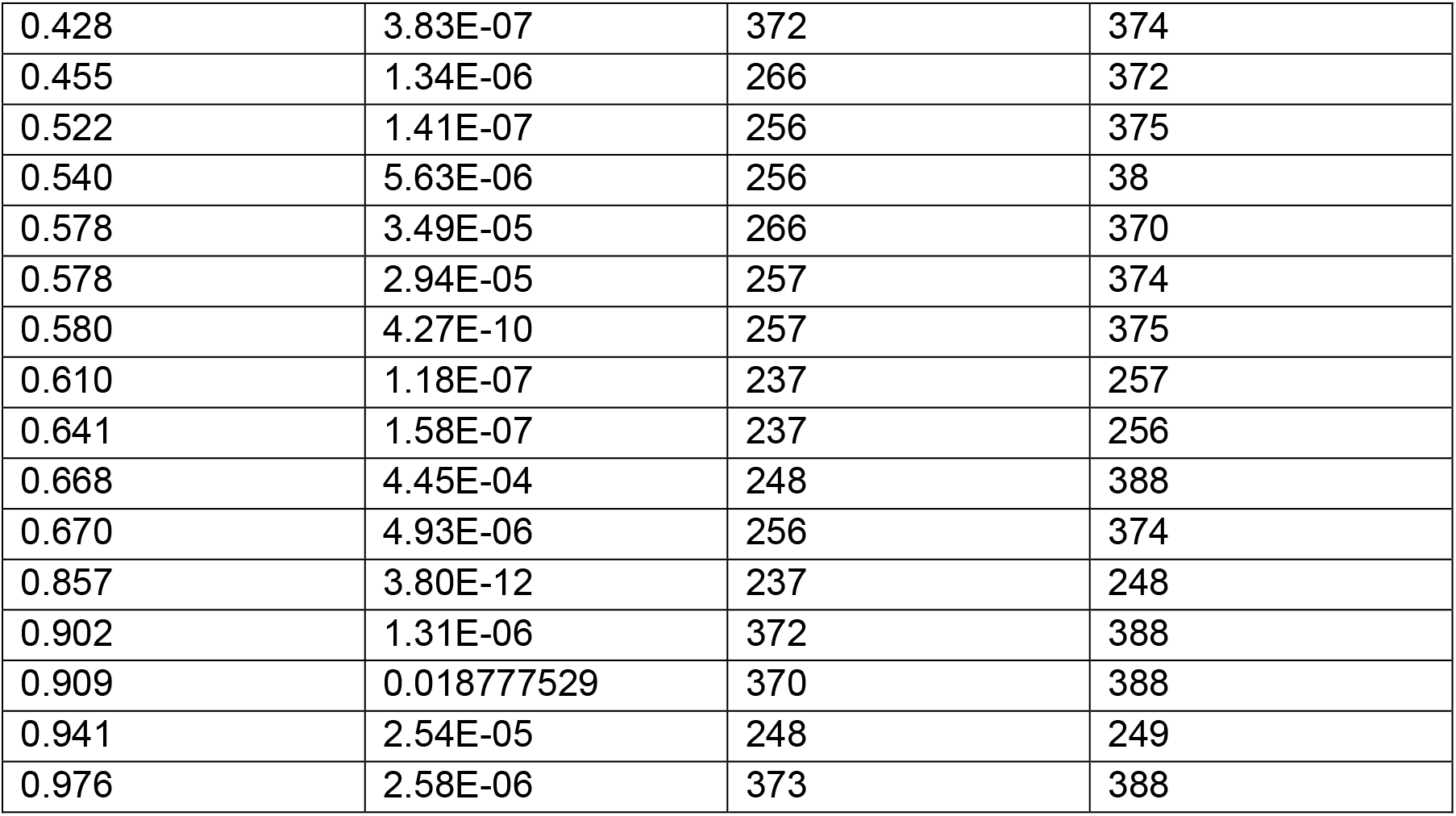
Crosslink fold changes with p-values less than 0.05 upon phase separation. The crosslink (xlink) position indicates the amino acid number of the N protein sequence. 169 in Xlink position 1 represents the K169-K65 crosslink. 169(phosphoS176) represents the phosphorylated form of this peptide, which is phosphorylated at S176.

| Fold change (log<sub>2</sub>) | p-value | Xlink position 1 | Xlink position 2 |
| --- | --- | --- | --- |
| -2.441 | 1.05E-04 | 257 | 266 |
| -2.226 | 5.88E-06 | 169 | 61 |
| -2.026 | 8.84E-08 | 266 | 347 |
| -1.697 | 1.23E-09 | 169 | 65 |
| -1.679 | 1.32E-09 | 169(phosphoS176) | 65 |
| -1.409 | 1.21E-05 | 38 | 61 |
| -1.324 | 3.20E-05 | 169 | 266 |
| -1.079 | 6.84E-04 | 169 | 38 |
| -0.941 | 3.61E-12 | 266 | 299 |
| -0.796 | 8.00E-07 | 102 | 61 |
| -0.765 | 2.94E-06 | 266 | 266 |
| -0.762 | 3.87E-04 | 342 | 38 |
| -0.730 | 5.18E-07 | 233 | 266 |
| -0.729 | 8.10E-05 | 266 | 38 |
| -0.715 | 0.003417897 | 355 | 375 |
| -0.648 | 0.008659112 | 102 | 266 |
| -0.607 | 0.001310878 | 342 | 355 |
| -0.594 | 2.84E-05 | 347 | 38 |
| -0.488 | 0.030217643 | 102 | 38 |
| -0.450 | 8.76E-04 | 266 | 375 |
| -0.448 | 1.53E-13 | 299 | 355 |
| -0.415 | 3.30E-06 | 233 | 38 |
| -0.382 | 0.048322087 | 38 | 65 |
| -0.380 | 2.21E-04 | 237 | 266 |
| -0.351 | 4.41E-04 | 266 | 388 |
| -0.277 | 0.0366727 | 266 | 374 |
| -0.264 | 2.72E-04 | 233 | 233 |
| -0.249 | 0.004338941 | 375 | 38 |
| -0.222 | 0.015033594 | 233 | 248 |
| -0.220 | 0.034461117 | 347 | 388 |
| -0.185 | 0.041924355 | 342 | 388 |
| 0.103 | 0.038548375 | 233 | 249 |
| 0.271 | 0.002666995 | 370 | 375 |
| 0.291 | 6.17E-06 | 233 | 237 |
| 0.347 | 6.53E-05 | 373 | 375 |
| 0.361 | 3.17E-06 | 237 | 388 |
| 0.378 | 5.90E-09 | 248 | 256 |
| 0.396 | 5.40E-05 | 256 | 342 |
| 0.398 | 1.41E-05 | 266 | 373 |
| 0.402 | 1.40E-06 | 257 | 342 |
| 0.402 | 9.55E-12 | 372 | 375 |
| 0.403 | 0.00139739 | 237 | 249 |
| 0.408 | 1.05E-07 | 248 | 257 |
| 0.418 | 0.002486181 | 370 | 374 |
| 0.428 | 3.83E-07 | 372 | 374 |
| 0.455 | 1.34E-06 | 266 | 372 |
| 0.522 | 1.41E-07 | 256 | 375 |
| 0.540 | 5.63E-06 | 256 | 38 |
| 0.578 | 3.49E-05 | 266 | 370 |
| 0.578 | 2.94E-05 | 257 | 374 |
| 0.580 | 4.27E-10 | 257 | 375 |
| 0.610 | 1.18E-07 | 237 | 257 |
| 0.641 | 1.58E-07 | 237 | 256 |
| 0.668 | 4.45E-04 | 248 | 388 |
| 0.670 | 4.93E-06 | 256 | 374 |
| 0.857 | 3.80E-12 | 237 | 248 |
| 0.902 | 1.31E-06 | 372 | 388 |
| 0.909 | 0.018777529 | 370 | 388 |
| 0.941 | 2.54E-05 | 248 | 249 |
| 0.976 | 2.58E-06 | 373 | 388 |

**Table S2.** A compilation of the SARS-CoV2 N protein phosphosites identified by mass spectrometry. **A.** The coverage map and phosphorylation sites for the N protein detected in proteomics experiments. Text in red indicates peptide regions that were not detected; green highlight indicates an unambiguous phosphorylation site; yellow highlight indicates an ambiguous phosphorylation site. **B.** Each site identified by our study is characterized as being unambiguously or ambiguously localized based on manual inspection of the product ion series. See Methods for the full protein sequence and a link to supporting evidence. S176 is a phosphorylation site identified on a crosslinked peptide.

| Phosphosite | Found in | Site Localization (this study) |
| --- | --- | --- |
| S21 | This study | Unambiguous |
| S23 | This study, | Unambiguous |
| T24 | This study, | Unambiguous |
| S26 | This study, (57) | Ambiguous (S26\|S33) |
| S33 | This study | Ambiguous (S26\|S33) |
| T76 | (10, 57) |  |
| S78 | This study, (10) | Ambiguous (S78\|S79) |
| S79 | This study, (10, 57) | Unambiguous |
| S105 | (10, 57) |  |
| T141 | (10) |  |
| T166 | (10) |  |
| S176 | This study, (10, 57) | Unambiguous |
| S180 | (10, 57) |  |
| S183 | (10, 57) |  |
| S184 | (10, 57) |  |
| S194 | (10, 57) |  |
| S197 | (57) |  |
| T198 | (10, 57) |  |
| S201 | (10, 57) |  |
| S202 | (10, 57) |  |
| T205 | (10, 57) |  |
| S206 | (10, 57) |  |
| T391 | (10) |  |
| T393 | (58) |  |
| S412 | This study | Ambiguous (S412\|S413) |
| S413 | This study | Ambiguous (S412\|S413) |
| S441 | This study | Unambiguous |

**Table S3.** Proteins identified by mass spectrometry in the high salt sample of N protein from human cells. **A.** The proteins were ranked by spectral abundance factor (SAF). These protein sequences were included in the CLMS search. Asterisks indicate exogenous proteins. **B.** Crosslinks between N and the G3BP1 and G3BP2 stress granule proteins identified in the qualitative CLMS experiment ranked by a support vector machine (SVM) score. Values are frequency of detection (two conditions with two replicates each). Xlink position indicates the amino acid number of the given protein sequence. SVM indicates the confidence that the crosslink is correctly identified.

| Gene | Unique Peptides | Percent Coverage | SAF | Species | UniProt Accession | Protein Name |
| --- | --- | --- | --- | --- | --- | --- |
| N | 57 | 70.0 | 104.6 | SARS2 | P0DTC9 | Nucleoprotein |
| HSPA1A | 54 | 59.3 | 27.5 | HUMAN | P0DMV8 | Heat shock 70 kDa protein 1A |
| G3BP2 | 38 | 42.5 | 19.5 | HUMAN | Q9UN86 | Ras GTPase-activating protein-binding protein 2 |
| G3BP1 | 37 | 60.5 | 17.8 | HUMAN | Q13283 | Ras GTPase-activating protein-binding protein 1 |
| HSPA8 | 27 | 37.8 | 9.8 | HUMAN | P11142 | Heat shock cognate 71 kDa protein |
| RPS12 | 2 | 17.4 | 8.3 | HUMAN | P25398 | 40S ribosomal protein S12 |
| NA | 4 | 17.3 | 7.4 | PIG* | P00761 | Trypsin |
| TUBB | 20 | 56.3 | 7.0 | HUMAN | P07437 | Tubulin beta chain |
| TUBA1B | 19 | 54.3 | 6.7 | HUMAN | P68363 | Tubulin alpha-1B chain |
| TUBB4B | 18 | 50.6 | 6.5 | HUMAN | P68371 | Tubulin beta-4B chain |
| KRT9 | 14 | 52.6 | 6.1 | HUMAN | P35527 | Keratin, type I cytoskeletal 9 |
| A2M | 27 | 20.7 | 5.8 | BOVIN* | Q7SIH1 | Alpha-2-macroglobulin |
| HSPA5 | 17 | 29.5 | 5.2 | HUMAN | P11021 | Endoplasmic reticulum chaperone BiP |
| NPM1 | 6 | 37.1 | 5.1 | HUMAN | P06748 | Nucleophosmin |

**Table S3. Proteins identified by mass spectrometry in the high salt sample of N protein from human cells.**
| Protein 1 | Xlink position 1 | Protein 2 | Xlink position 2 | SVM score |
| --- | --- | --- | --- | --- |
| G3BP2_HUMAN | 50 | G3BP2_HUMAN | 281 | 2.17 |
| Nucleoprotein | 375 | G3BP1_HUMAN | 453 | 1.63 |
| G3BP2_HUMAN | 50 | Nucleoprotein | 266 | 1.56 |
| G3BP2_HUMAN | 50 | Nucleoprotein | 102 | 1.26 |
| G3BP2_HUMAN | 370 | G3BP2_HUMAN | 407 | 1.04 |
| G3BP2_HUMAN | 365 | G3BP2_HUMAN | 407 | 0.88 |
| G3BP2_HUMAN | 50 | Nucleoprotein | 38 | 0.71 |
| Nucleoprotein | 388 | G3BP1_HUMAN | 453 | 0.67 |
| Nucleoprotein | 266 | G3BP1_HUMAN | 393 | 0.14 |
| Nucleoprotein | 374 | G3BP1_HUMAN | 453 | 0.00 |

**Table S4.** Quantitative analysis of N condensates. The parameters of fitting to a dose-response equation in the presence and absence of polyC RNA in response to increasing concentrations of salt or nelfinavir mesylate.

| Condition | IC<sub>50</sub> / EC<sub>50</sub> | *n* | R<sup>2</sup> | Figure |
| --- | --- | --- | --- | --- |
| N with NaCl, condensate volume | 90 ± 1 mM | 12.26 ± 0.06 | 1 | 1D |
| N + PolyC with NaCl, condensate volume | 152 ± 6 mM | 11 ± 9 | 0.97 | 1D |
| N + PolyC with nelfinavir mesylate, condensate number | 4.3 ± 1.3 μM | 1.9 ± 1.0 | 0.90 | 5B |

**Figure 1_figure supplement 1.**
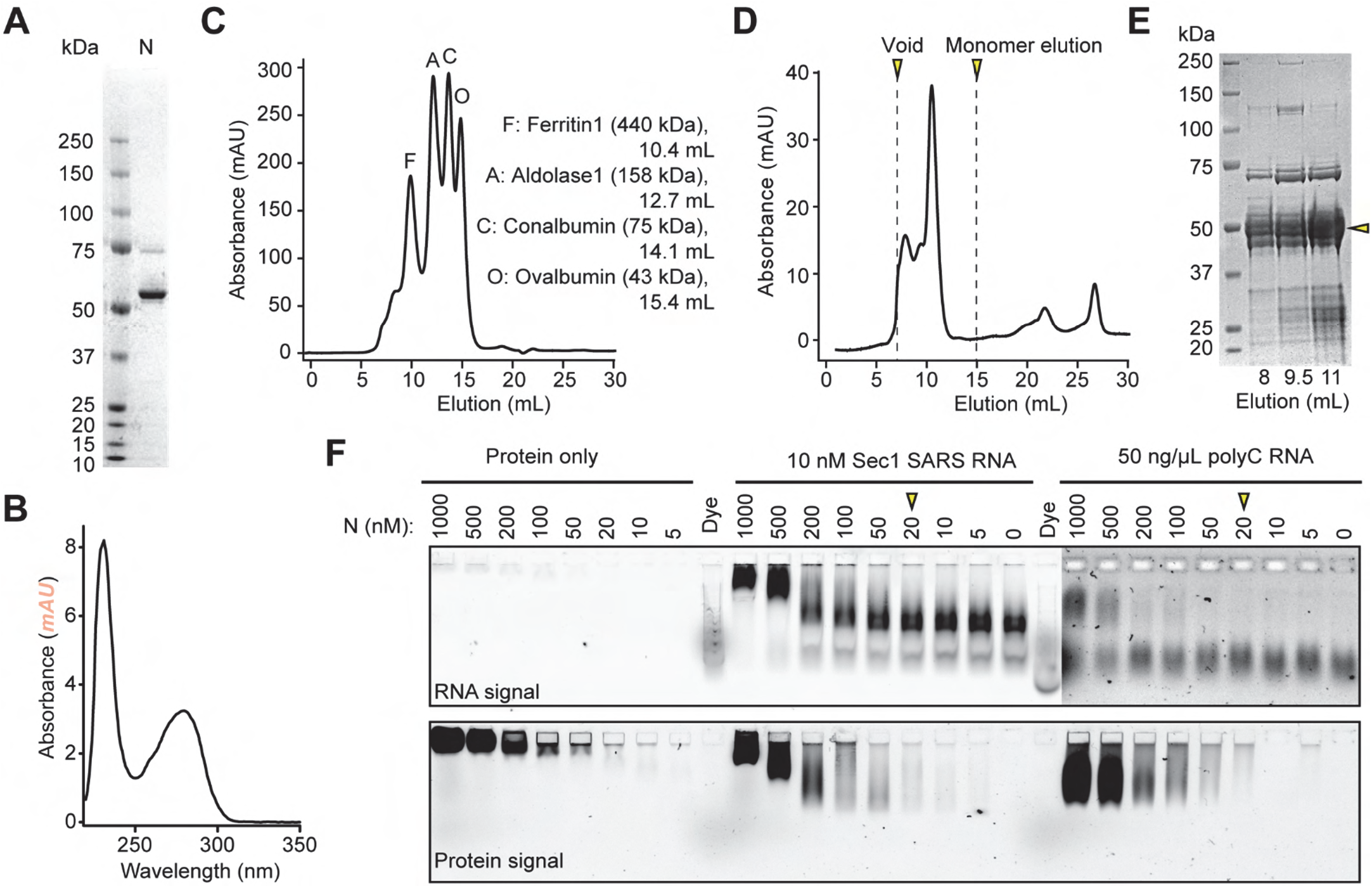
Gel filtration and EMSA analysis of the WT N protein purified from HEK293S GNTI-cells. **(A)** The coomassie-stained denaturing gel of the N protein purified from affinity chromatography. **(B)** UV absorbance of the N protein purified from affinity chromatography shows no evidence for the presence of contaminating nucleic acids. **(C)** UV absorbance of protein standards eluting from a gel filtration column. **(D)** UV absorbance of the N protein eluting from a gel filtration column. Arrows mark the void volume and expected elution volume for an N monomer. **(E)** The coomassie-stained denaturing gel of the eluents from the gel filtration column. **(F)** EMSA using no RNA, 10 nM Section 1 viral RNA, and 50 ng/μl polyC RNA and decreasing concentration of N protein. The protein was labeled with LD655. RNA was labeled with Cy3. Arrows indicate the minimum protein concentration for each condition with a noticeable shift in the protein band.

**Figure 1_figure supplement 2.**
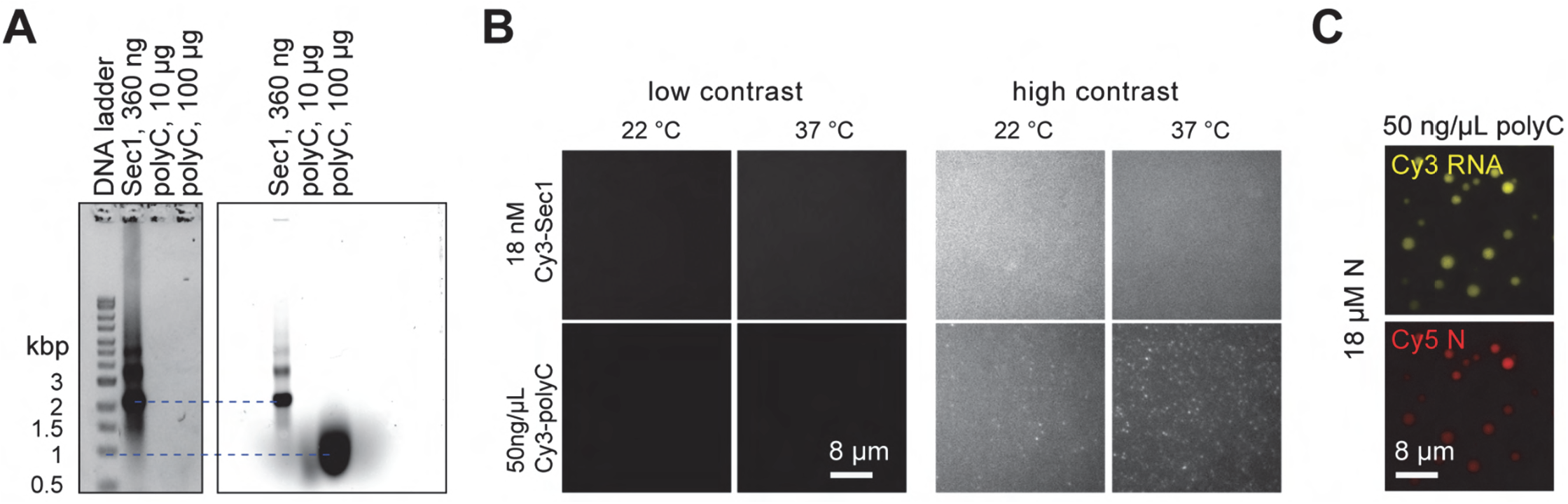
RNA substrates do not form condensates in the absence of N protein. (**A**) An agarose gel picture of Section 1 viral RNA and polyC RNA substrates. The gel was stained with GelRed (left) and the RNA substrates were labeled with Cy3 (right). The ladder corresponds to the length of double-stranded DNA. Estimated lengths of Section 1 viral RNA and polyC RNA are 6 kb and 2 kb, respectively. (**B**) Representative pictures show that 18 nM Section 1 and 50 ng/μl polyC RNA do not form condensates in the absence of N protein. The assay was performed in 150 mM NaCl. (**C**) Two-color imaging shows colocalization of LD655-labeled N protein and Cy3-labeled polyC RNA in condensates.

**Figure 1_figure supplement 3.**
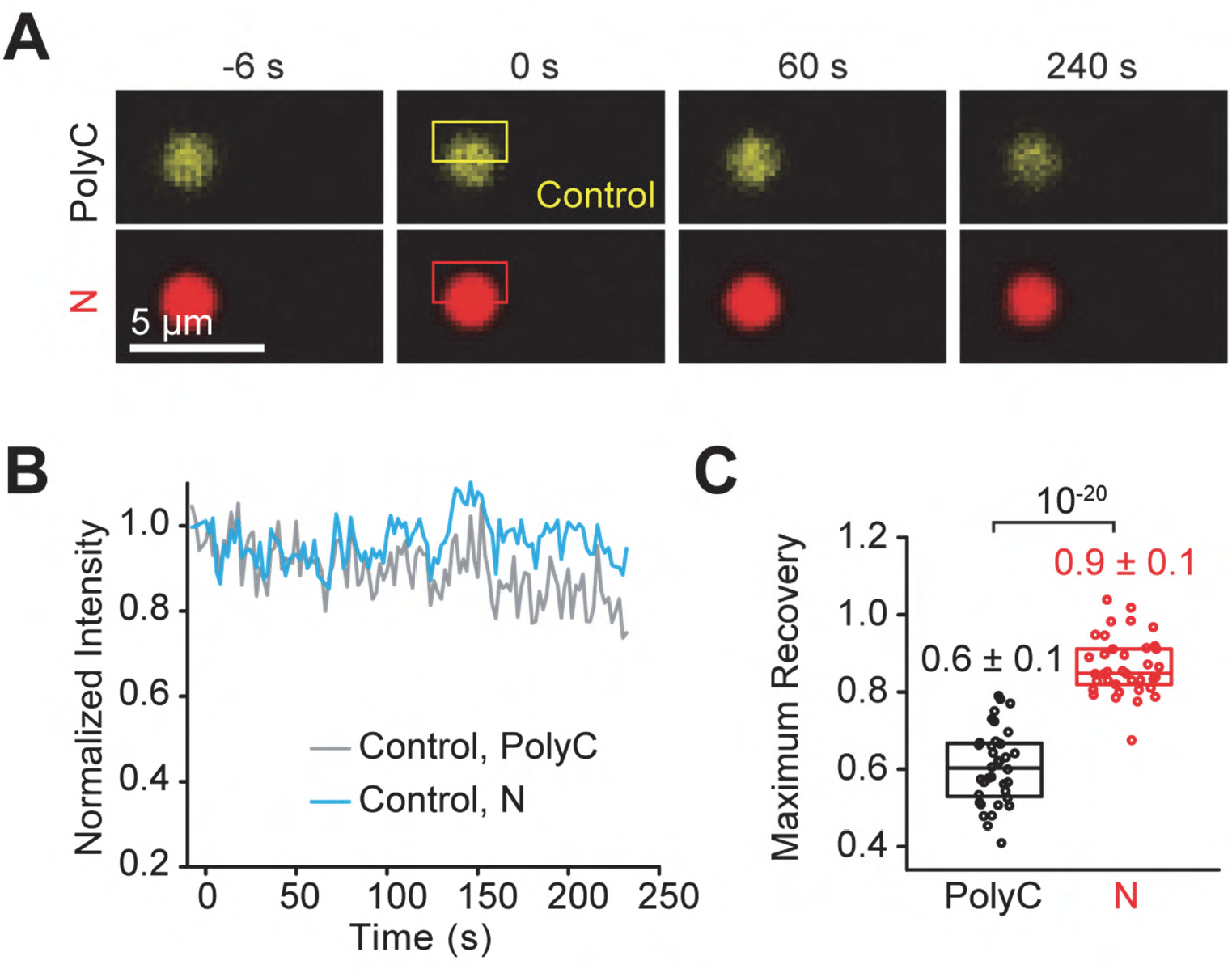
FRAP analysis of N-polyC condensates in vitro. **(A)** Control experiments show changes in the fluorescent signal of N and polyC without photobleaching. **(B)** The changes in the integrated fluorescent intensities of regions highlighted with yellow and red rectangles in A. **(C)** The maximum fractional recovery of N and polyC in condensates after photobleaching (n =37, mean ± s.d.). The center and edges of the box represent the median with the first and third quartiles. The p-value was calculated from a two-tailed t-test.

**Figure 2_figure supplement 1.**
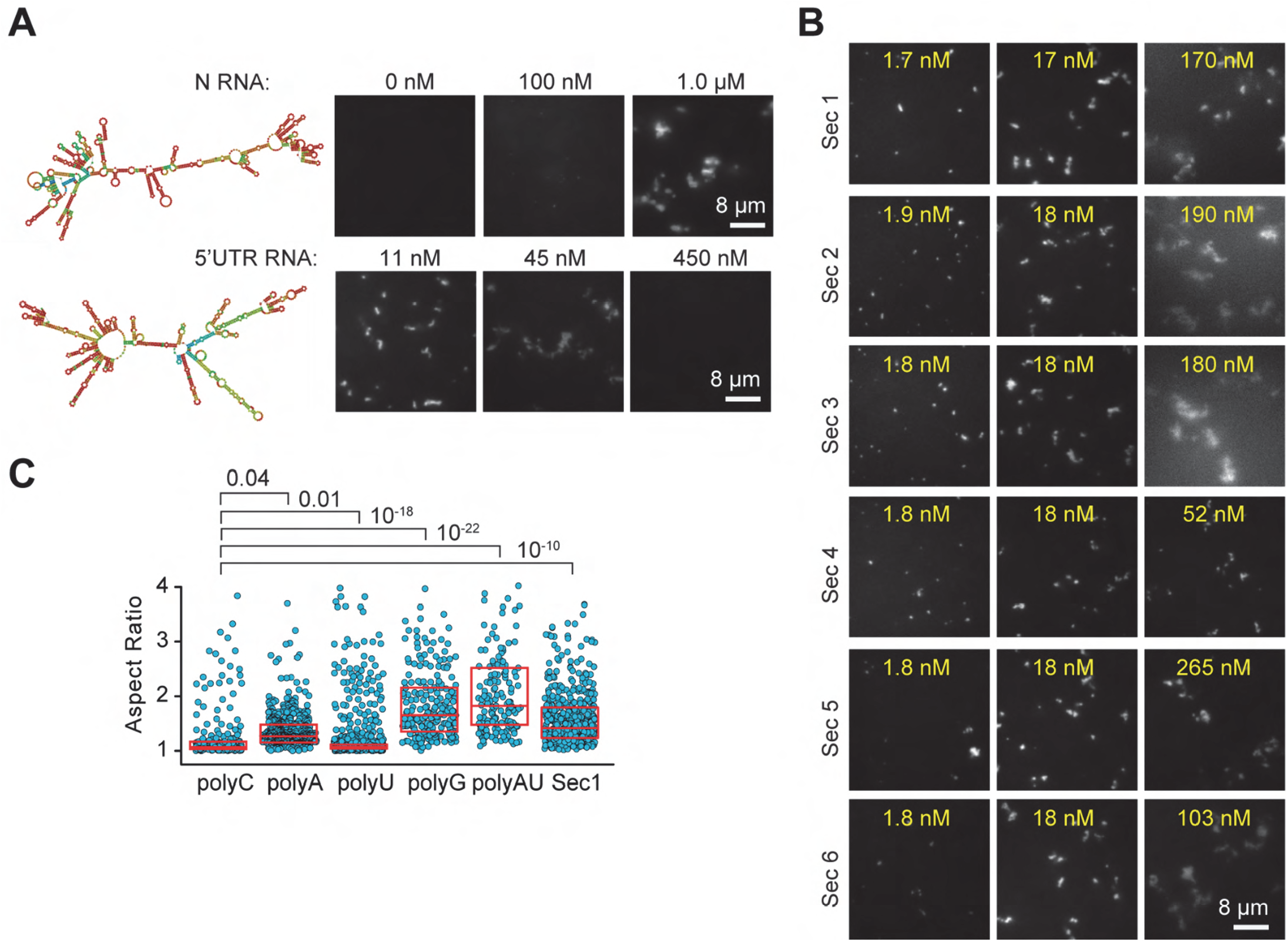
The N protein forms asymmetric condensates with viral RNA. **(A)** Structure prediction of viral RNA (left) and the formation of asymmetric N condensates under different RNA concentrations (right). The N RNA is the 1.3 kb long genomic RNA fragment that encodes the SARS CoV-2 N protein. The 5’ UTR RNA is the first 1,000 bases of the SARS-CoV-2 genome. The N protein concentration was set to 18.5 μM (IVT: in vitro transcribed; UTR: untranslated region). **(B)** The formation of asymmetric N condensates under different viral RNA concentrations. The N protein concentration was set to 18.5 μM. **(C)** The distribution of aspect ratios of individual condensates formed with different RNA substrates. The N protein concentration was set to 18.5 μM, and RNA concentration was set to 50 ng/μL for RNA homopolymers and 18 nM for Sec1 RNA. The center and edges of the box represent the median with the first and third quartiles. P values are calculated from two-tailed t-tests.

**Figure 2_figure supplement 2.**
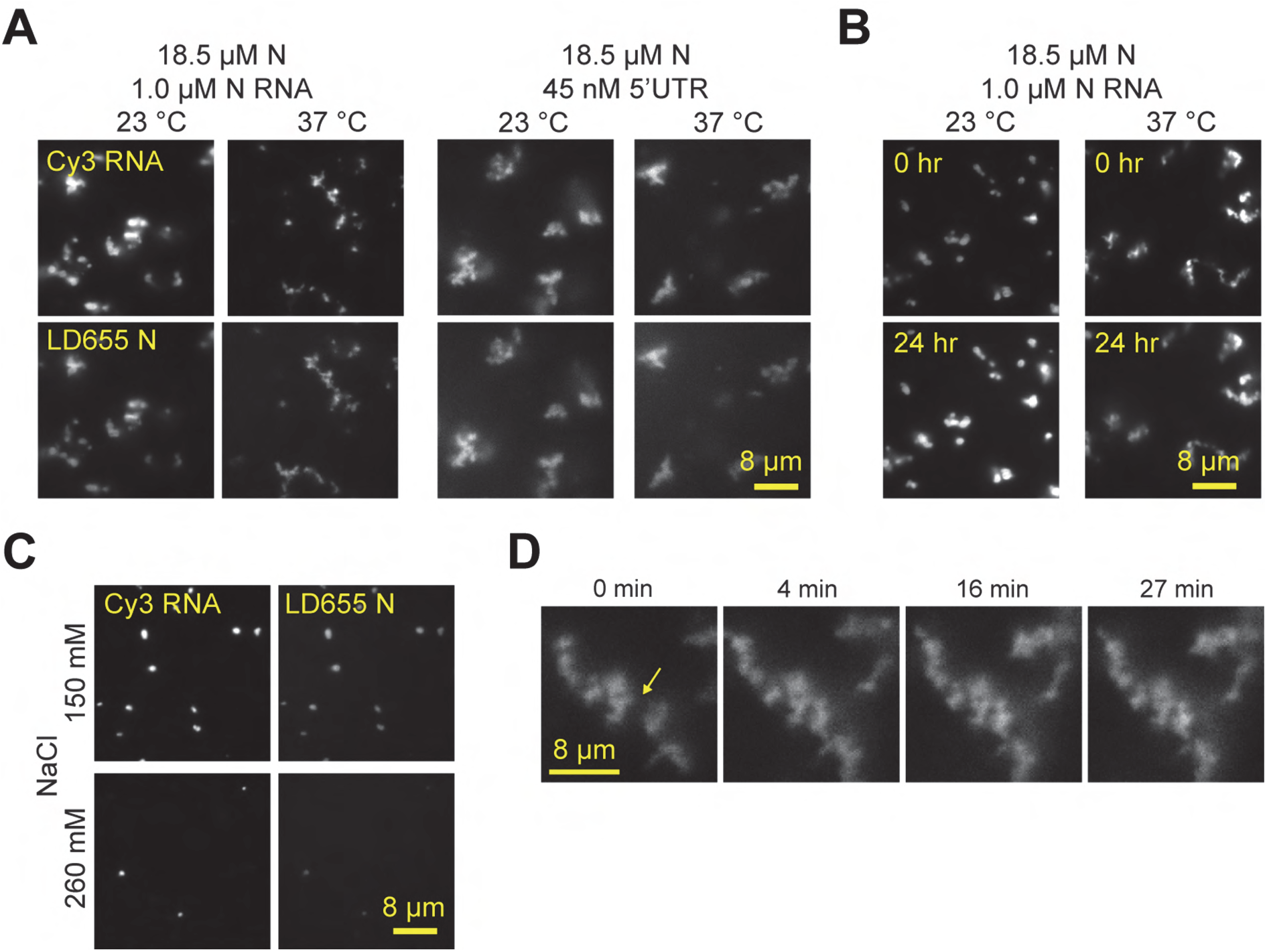
Condensates formed by the N protein and viral RNA do not fuse or change shape but are dissolved at higher salt. **(A)** Condensates formed by the N protein with in vitro transcribed N RNA or the 5’UTR of the viral RNA are not sensitive to an increase of temperature to 37 °C. **(B)** Condensates formed by the N protein with 5’UTR of viral RNA do not change shape over 24 hours. **(C)** Condensates formed by 11.5 μM N protein and 36 nM 5’UTR RNA are dissolved in the presence of 260 mM NaCl. **(D)** Condensates formed by 18.5 μM N protein and 18 nM Sec1 RNA do not fuse after contact (yellow arrow).

**Figure 2_figure supplement 3.**
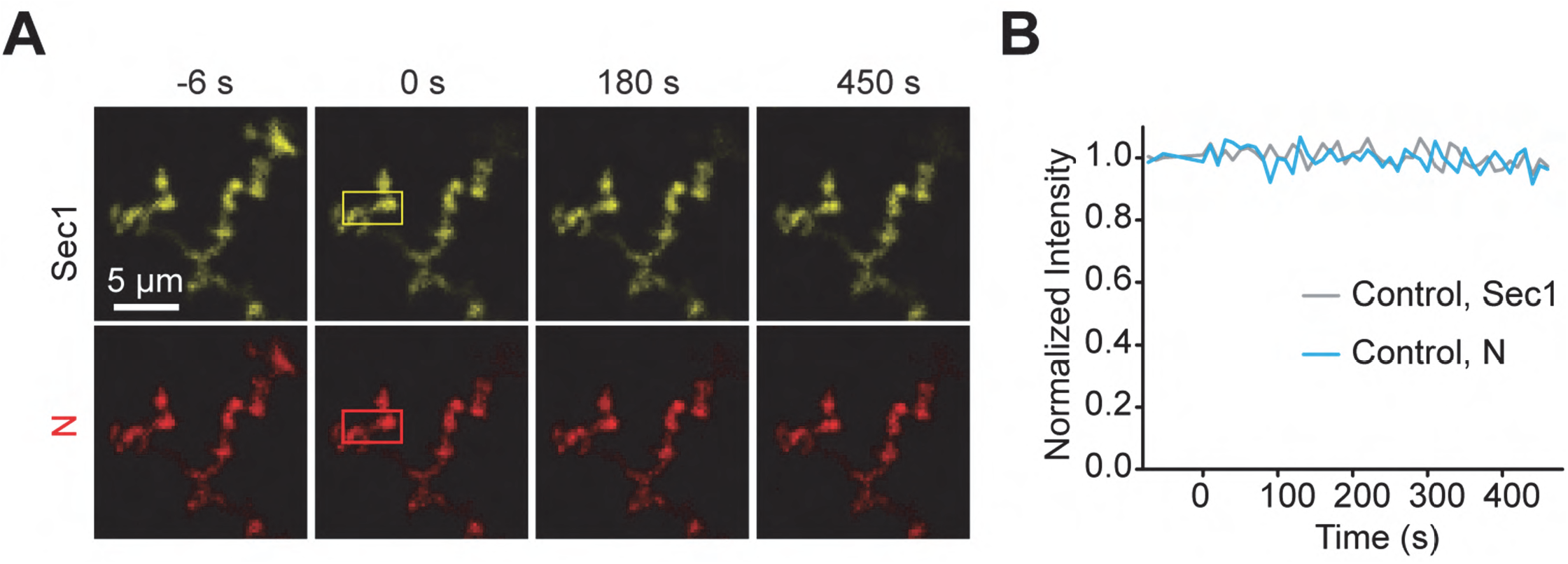
FRAP analysis of N-Sec1 condensates in vitro. (**A**) Control experiments show snapshots of N and Sec1 without photobleaching. (**B**) The changes in the integrated fluorescent intensities of regions highlighted with yellow and red rectangles in A.

**Figure 3_figure supplement 1.**
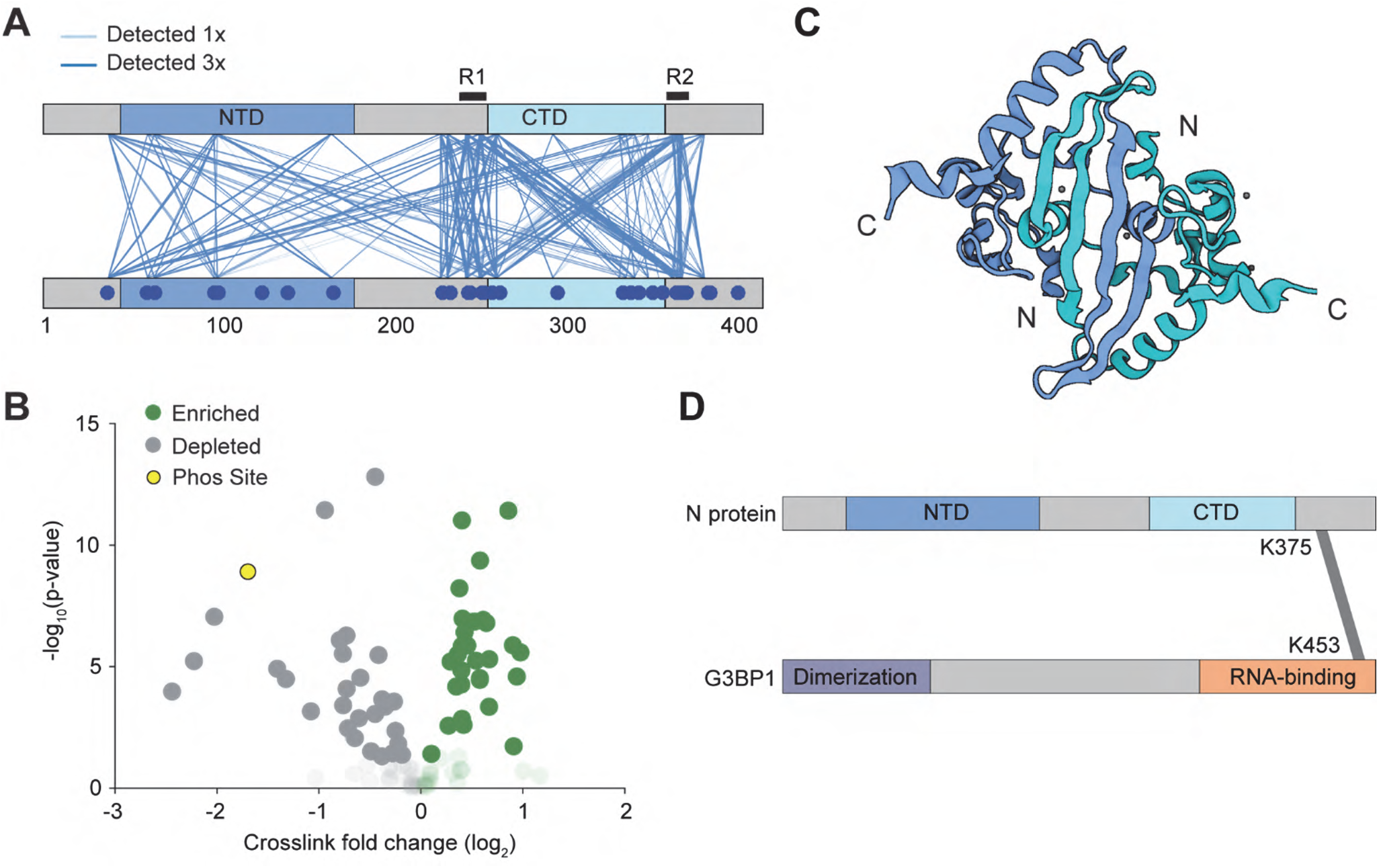
Interactions between N proteins and between N and G3BP1. **(A)** In an initial qualitative experiment, pairwise interactions were detected by CLMS at 300 mM salt. Lines depict a unique crosslink detected. The regions of N protein interactions flank CTD. **(B)** Volcano plot of the quantitative CLMS data comparing the condensate and no condensate condition. Opaque data points have a p-value below 0.05, transparent data points have a p-value greater than 0.05. Green represents unique crosslinks that are enriched in the condensate condition. The yellow markers represent the K169-K65 and K169 (phosS176)-K65 crosslinks. **(C)** The structure of the CTD of the SARS-CoV-2 N protein was plotted with BioRender (PDB 6WJI (*78*)). Because the N and C termini of the protomers are positioned away from each other, R1 and R2 within the same dimer are unlikely to interact with each other. **(D)** Mass spectrometry identified that the RNA binding domain of the stress granule protein G3BP1 interacts with the R2 region of N.

**Figure 4_figure supplement 1.**
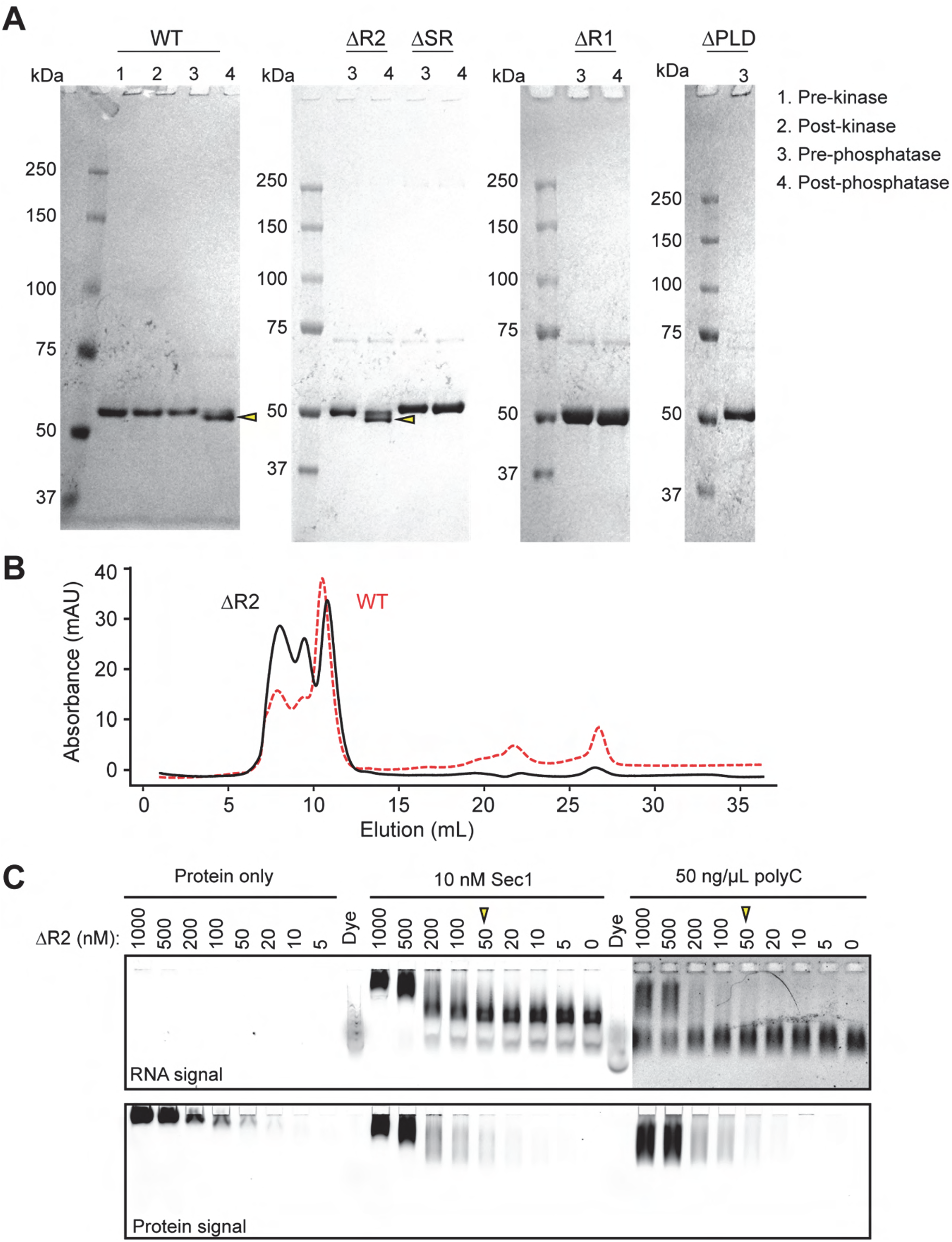
Purification and characterization of the deletion mutants. **(A)** Denaturing gel pictures of purified WT and deletion mutants of N protein in the presence and absence of kinase and phosphatase treatment (see Methods). The gels were stained with Coomassie. Yellow arrowheads highlight reduction in molecular weight upon treatment with λ phosphatase. (**B**) UV absorbance of the ΔR2 mutant eluting from a gel filtration column. UV absorbance of WT N under the same experimental conditions is shown in a dashed red curve for comparison. **(C)** EMSA gels using no RNA, 10 nM Section 1 viral RNA, or 50 ng/μl polyC RNA and decreasing concentration of the ΔR2 mutant. The protein was labeled with LD655. RNA was labeled with Cy3. Arrows indicate the minimum protein concentration for each condition with a noticeable shift in the protein band.

**Figure 4_figure supplement 2.**
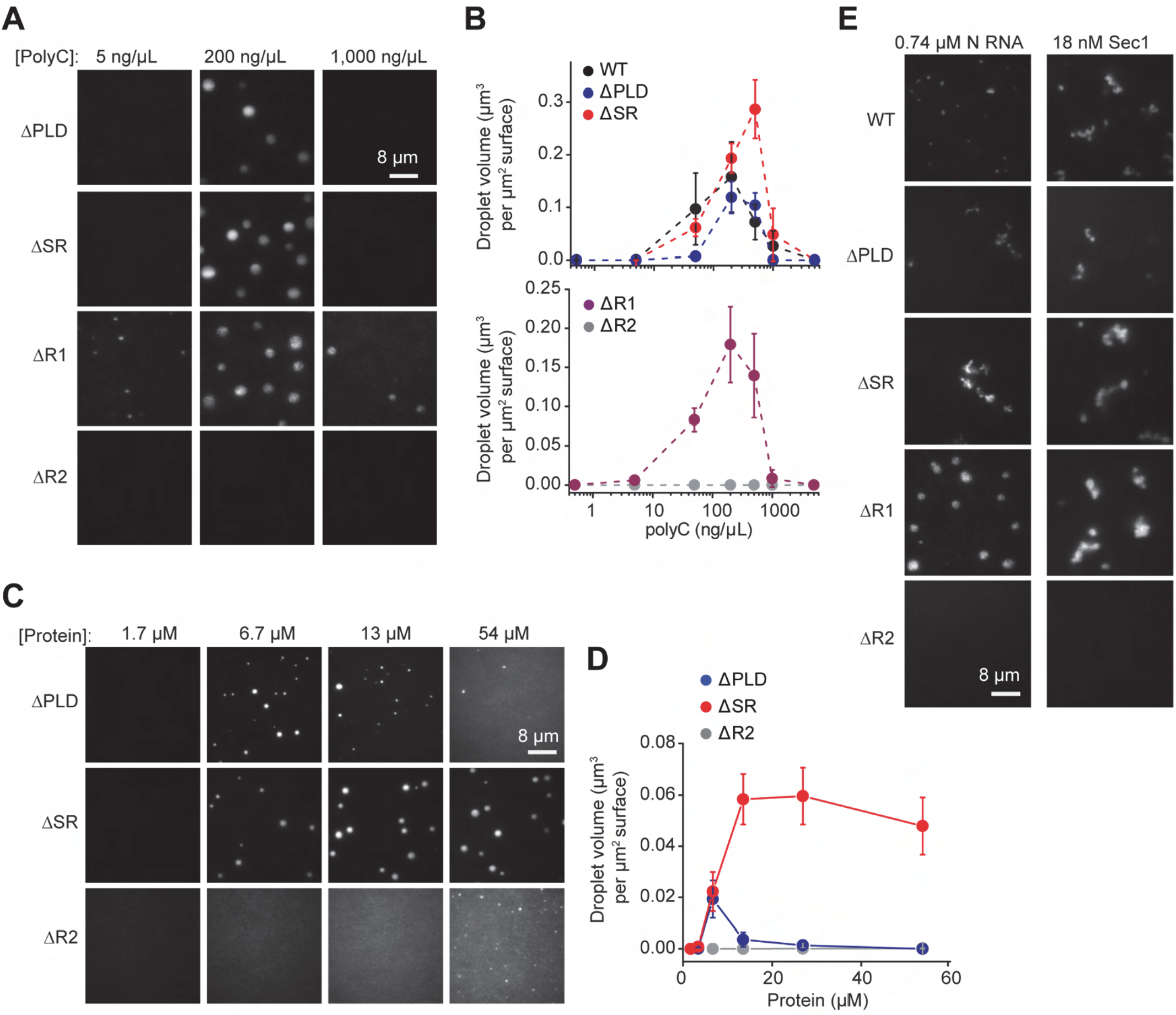
The in vitro phase of the truncation mutants of the N protein. **(A)** Example pictures show that the N truncation mutants, except ΔR2, form spherical condensates with polyC RNA under different RNA concentrations. The N protein concentration was set to 18.5 μM. **(B)** The total volume of N-RNA condensates settled per micron squared area on the coverslip (mean ± s.d.; n = 20, two technical replicates) exhibits a reentrant behavior under an increasing RNA concentration. **(C)** Example pictures show that the N truncation mutants, except ΔR2, form spherical condensates with polyC RNA under different protein concentrations. The polyC RNA concentration was set to 50 ng/μl. **(D)** The total volume of N-RNA condensates settled per micron squared area on the coverslip (mean ± s.d.; n = 20, two technical replicates) under an increasing protein concentration. **(E)** Phase separation of truncated N protein constructs with 0.74 μM in vitro transcribed N RNA or 18 nM Sec1 RNA. The protein concentration was set to 18.5 μM.

**Figure 4_figure supplement 3.**
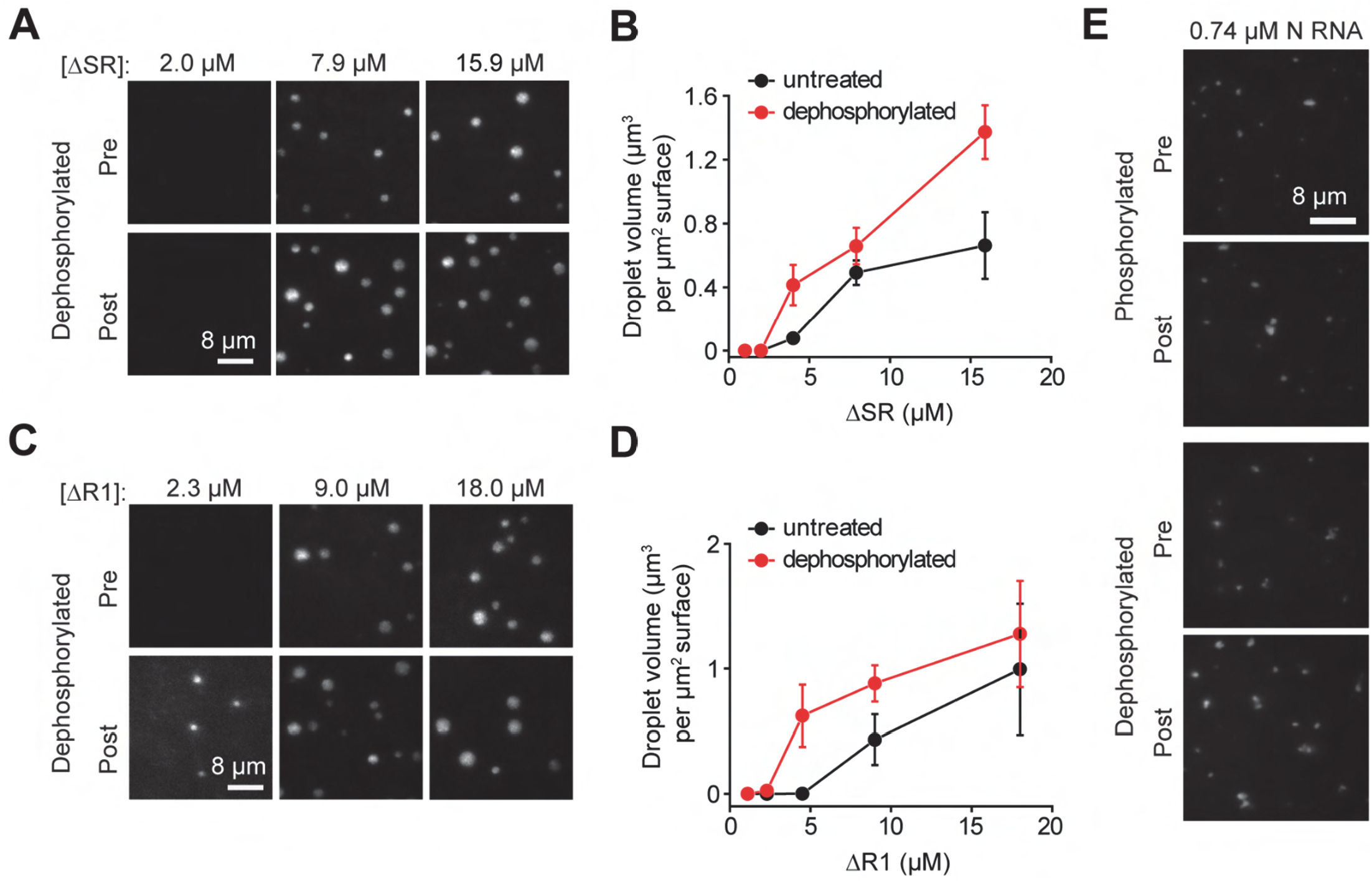
Phase separation of truncated N proteins under different phosphorylation conditions in vitro. **(A)** Images of condensates formed by untreated and dephosphorylated ΔSR in the presence of 50 ng/μL polyC RNA. **(B)** The total volume of the condensates settled per micron squared area on the coverslip as a function of ΔSR concentration (mean ± s.d., n = 20 with two technical replicates). **(C)** Images of condensates formed by untreated and dephosphorylated ΔR1 in the presence of 50 ng/μL polyC RNA. **(D)** The total volume of the condensates settled per micron squared area on the coverslip as a function of ΔR1 concentration (mean ± s.d., n = 20 with two technical replicates). **(E)** Full-length N protein forms asymmetric condensates with 0.74 μM in vitro transcribed N RNA before and after phosphorylation and dephosphorylation. The protein concentration was set to 35 μM.

**Figure 5_figure supplement 1.**
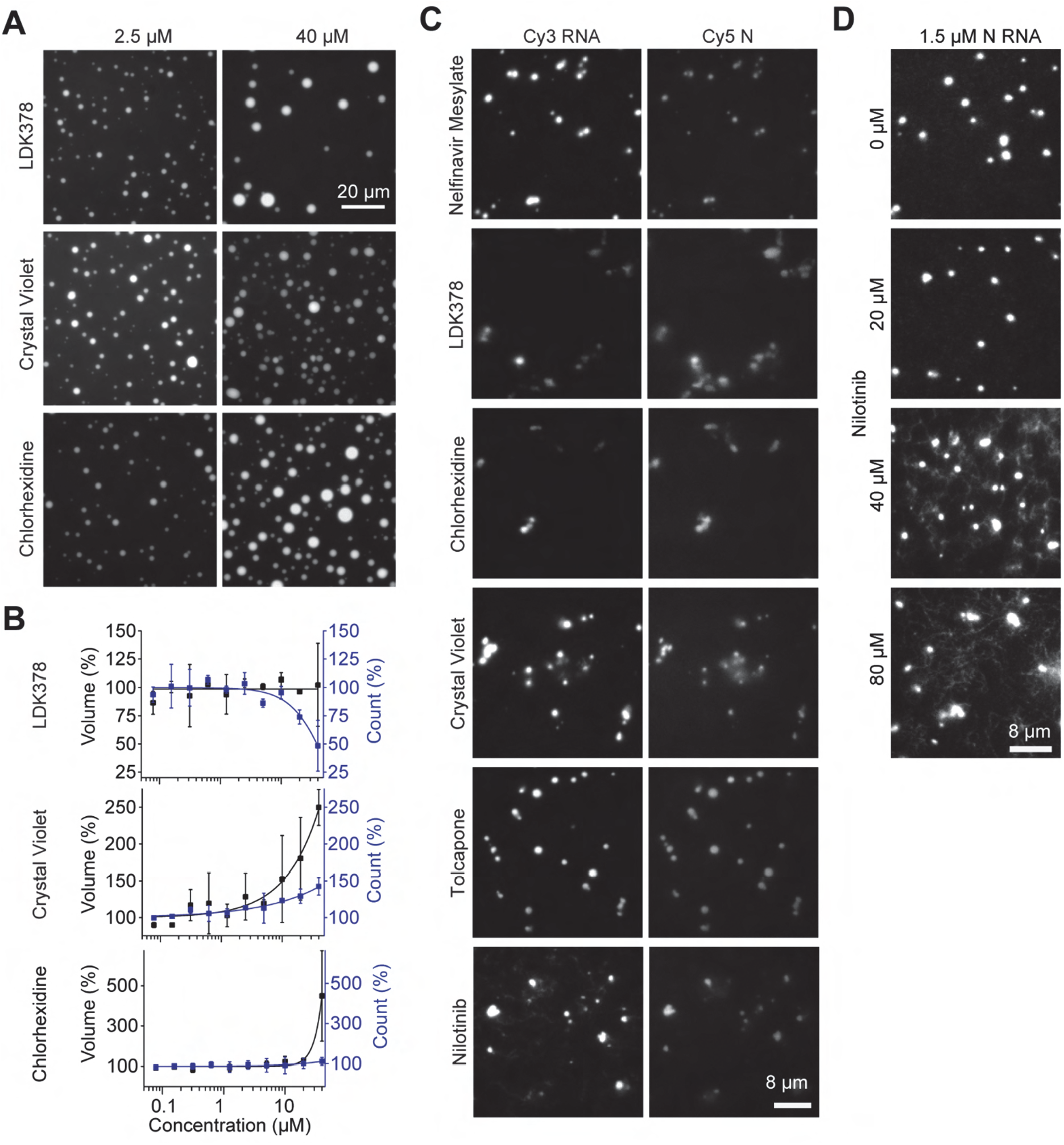
Phase separation of the N protein under drug treatment. **(A)** Example pictures show phase separation of 7.8 μM N protein and 50 ng/μL polyC RNA at different drug concentrations. **(B)** The percent change on the number (blue) and total volume (black) of N-polyC condensates settled per micron squared area on the coverslip under different drug concentrations (mean ± s.d., n = 8 with two technical replicates). **(C)** Example pictures show phase separation of 57.6 μM LD655-labeled N protein and 1.5 μM Cy3-labeled in vitro transcribed N RNA in the presence of different drug treatments. The concentration of drugs was set to 40 μM. **(D)** Condensates formed in the presence of 57.6 μM N protein and 1.5 μM N RNA form thread-like filaments at high nilotinib concentrations.

**Figure 5_figure supplement 2.**
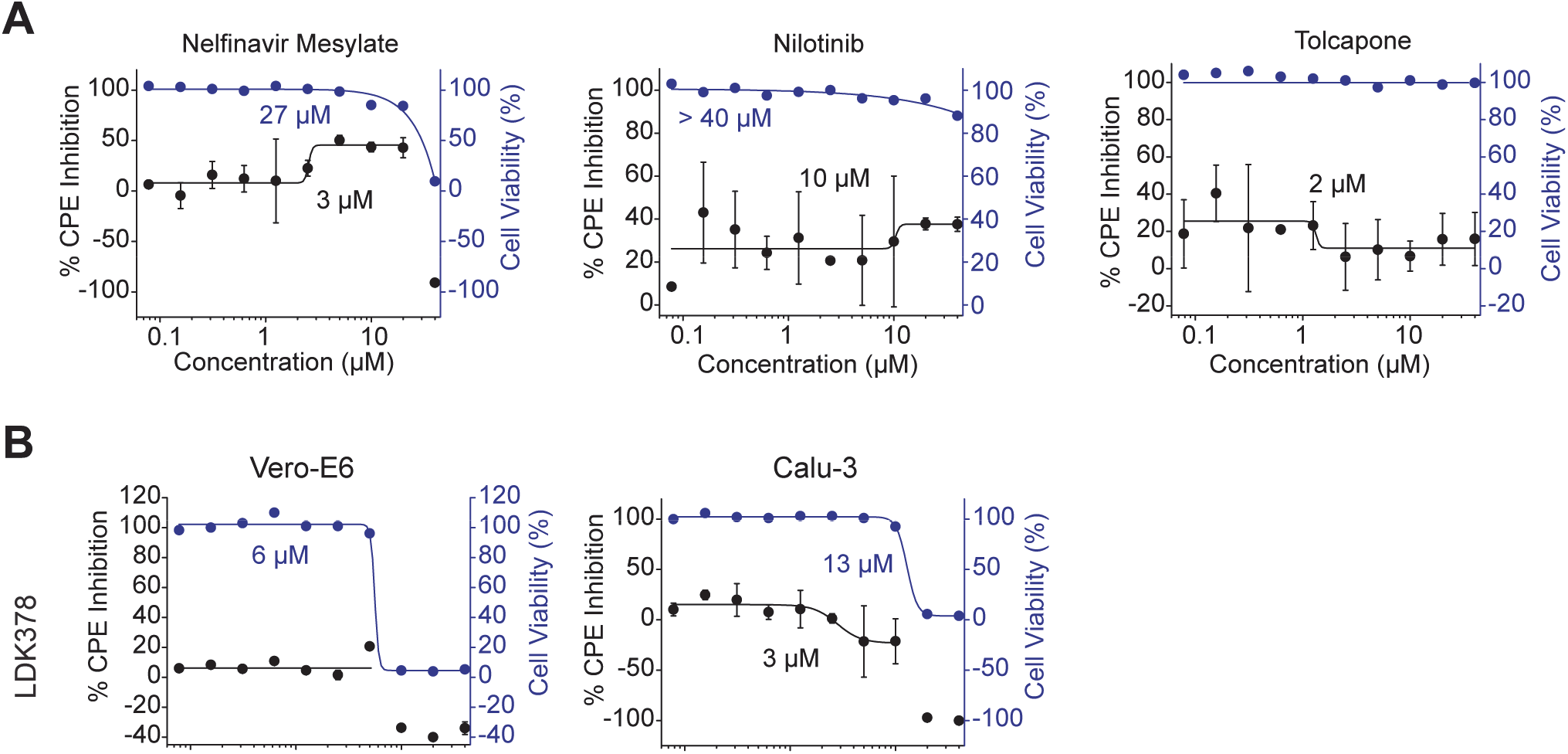
The viability of SARS-CoV-2 infected cell under drug treatment. **(A)** Percent CPE inhibition (black, mean ± s.d., two technical replicates) and cell viability (blue, mean) of SARS-CoV-2 infected Calu3 cells treated with serial dilutions of drugs. Solid curves represent a fit to a dose-response equation to determine EC_50_. **(B)** Percent CPE inhibition (black, mean ± s.d., two technical replicates) and cell viability (blue, mean) of SARS-CoV-2 infected Vero-E6 and Calu3 cells treated with serial dilutions of LDK378. Solid curves represent a fit to a dose-response equation to determine EC_50_ (Table S4).

**Figure 6_figure supplement 1.**
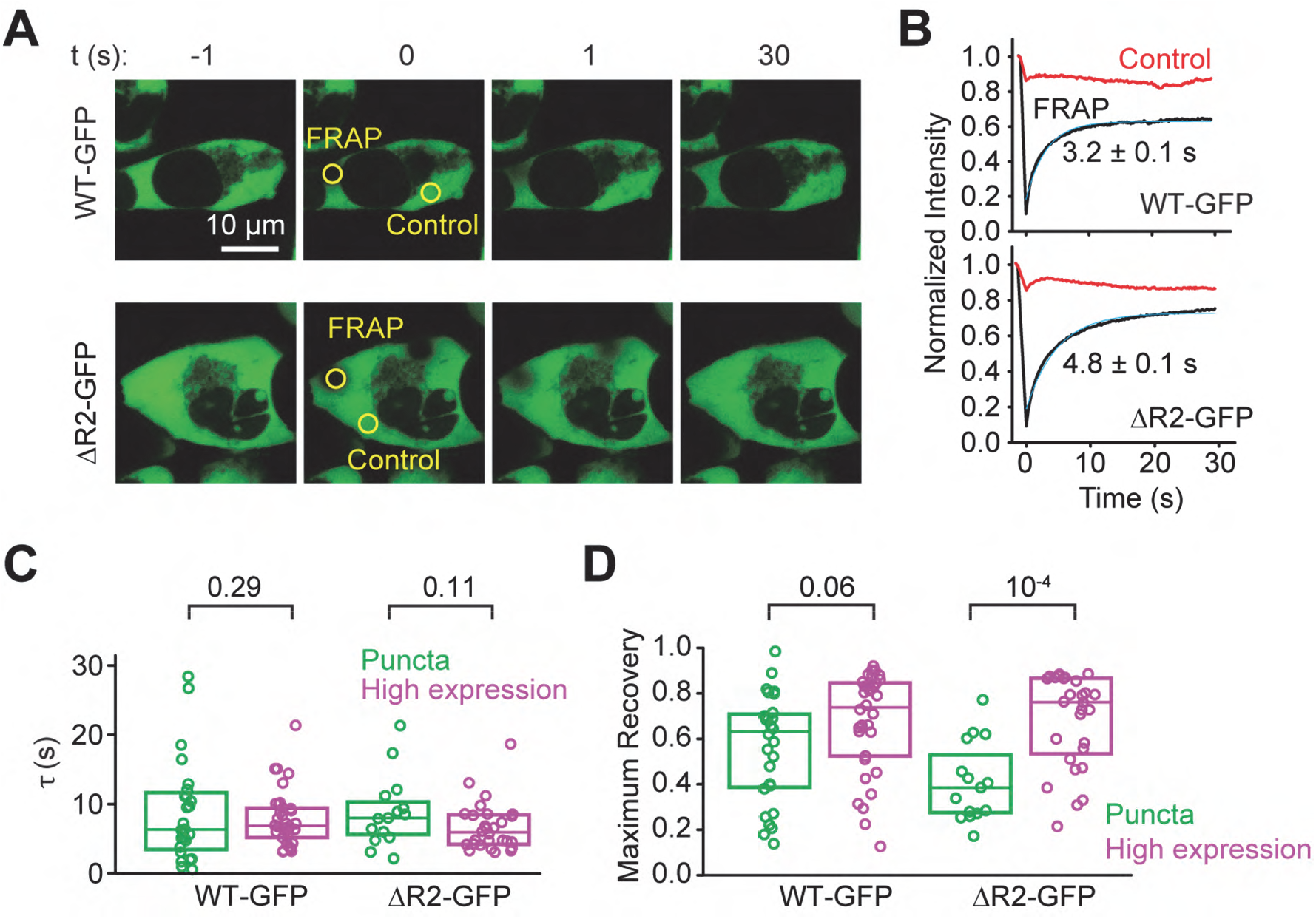
Dynamics of N protein in high-expressing cells. **(A)** Representative FRAP imaging of a cell with a high expression of WT or ΔR2 N. Circles show the photobleached and control (not bleached) regions. **(B)** Fluorescence recovery signals of the N protein in the bleached versus the control regions. The solid curve represents a single exponential fit to reveal the recovery lifetime (τ, ±95% confidence interval). **(C)** The distribution of fluorescence recovery lifetimes of cells expressing WT or ΔR2 N and exhibiting either puncta or high expression (from left to right, n=28, 34, 15, and 28). The center and edges of the box represent the median with the first and third quartiles. The p values were calculated from a two-tailed t-test. **(D)** The maximum fractional recovery after photobleaching of cells expressing WT or ΔR2 N and exhibiting either puncta or high expression (from left to right, n=28, 34, 15, and 28). The center and edges of the box represent the median with the first and third quartiles. The p values were calculated from a two-tailed t-test.

**Figure 6_figure supplement 2.**
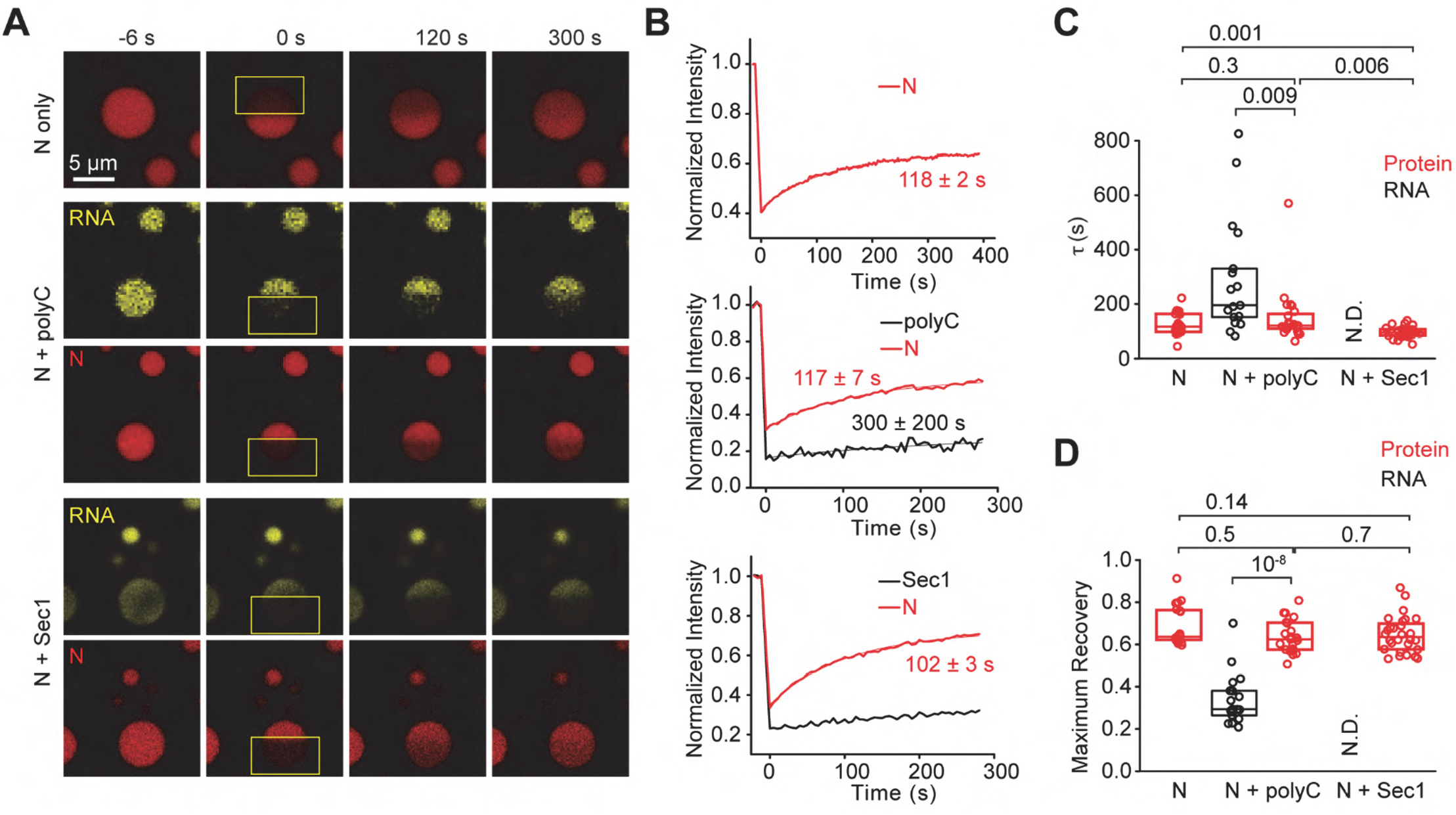
Dynamics of WT N protein in the presence of low (50 mM NaCl) salt. **(A)** Representative FRAP imaging of N protein only, or in the presence of polyC or Sec1 RNA. The concentrations of N, polyC, and Sec1 were kept at 24 μM, 50 ng/μL, and 18 nM, respectively. Rectangles highlight the photobleached area. **(B)** Fluorescence recovery signals of the N protein and RNA in the bleached versus the control regions. Solid curves represent a single exponential fit to reveal the recovery lifetime (τ, ±95% confidence interval). **(C)** The distribution of fluorescence recovery lifetimes of the condensates (from left to right, n = 20, 20, 23, 28, and 28). The center and edges of the box represent the median with the first and third quartiles. The p values were calculated from a two-tailed t-test. **(D)** The maximum fractional recovery after photobleaching (from left to right, n = 20, 20, 23, 28, and 28). The center and edges of the box represent the median with the first and third quartiles. The p values were calculated from a two-tailed t-test.

**Figure 6_figure supplement 3.**
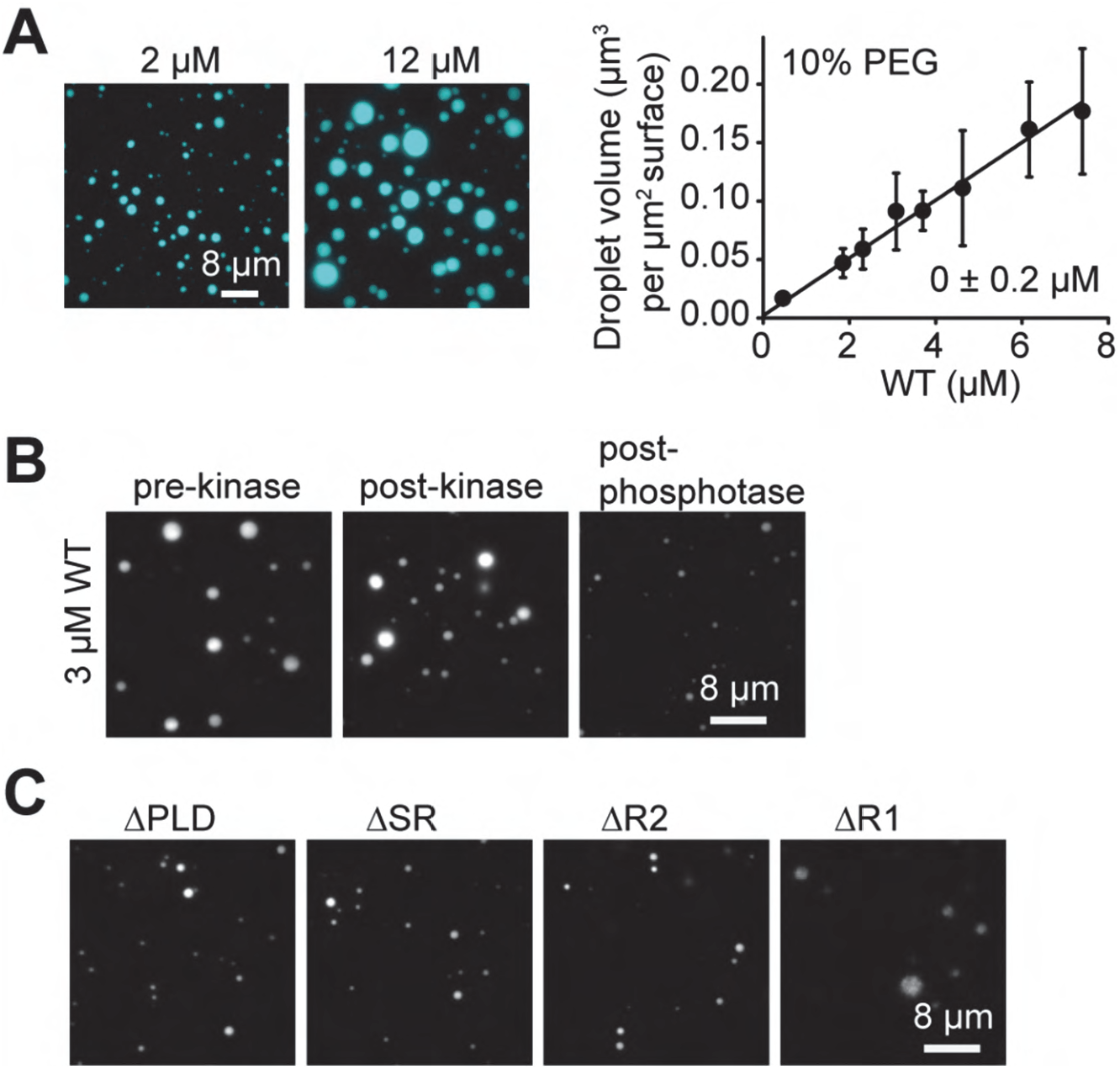
Phase separation of WT and truncation mutants in the presence of a crowding agent. **(A)** (Left) Images of the LD655-labeled N protein in the presence of 10% PEG and 150 mM NaCl. (Right) The total volume of N-RNA condensates settled per micron squared area on the coverslip (mean ± s.d., n = 20 with two technical replicates). A linear fit (solid line) reveals c_sat_ (± s.e.). **(B)** Images of the condensates formed in the presence of LD655-labeled WT N protein before or after phosphatase and kinase treatments in the presence of 10% PEG and 150 mM NaCl. **(C)** Images of the LD655-labeled truncated N protein mutants in the presence of 10% PEG and 150 mM NaCl. The protein concentration was set at 3 μM. Assays were performed in the absence of RNA.

## Methods

### Protein purification

A construct for expressing N protein with a C-terminal Strep-tag was obtained from the Krogan lab (UCSF). A C-terminal yBBr labeling site was added for labeling with a fluorescent dye. Truncation mutants, ΔPLD (amino acids 2-30: SDNGPQNQRNAPRITFGGPSDSTGSNQNG), ΔSR (amino acids 183-195, SSRSSSRSRNSSR), ΔR1 (amino acids 235-256, SGKGQQQQGQTVTKKSAAEASK) and ΔR2 (amino acids 369-390, KKDKKKKADETQALPQRQKKQQ) were generated using Gibson cloning (New England Biosciences). Sufficient amounts of DNA were obtained growing 1 L of transfected XL1 Blue *Escherichia coli* cells overnight and performing a gigaprep (Zymo). For protein expression, HEK293S GNTI- cells (RRID: CVCL_A785) were grown in suspension in Freestyle media (Gibco) supplemented with 2% FBS (VWR) and 1% penicillin-streptomycin (Gemini Bio-products) to 2 million cells/mL. Cells were spun down for 10 min at 1,200 *g* and resuspended in fresh, antibiotic-free media. For 250 mL cells, the transfection solution was created by mixing 1.8 mL PEI (1mg/mL, pH 7.0 in PBS) dissolved in 20 mL Freestyle media and 0.66 mg DNA dissolved in 20 mL Freestyle media. The mixture was incubated at room temperature for 15 min before being adding it to the cell culture. Transfected cell culture was grown for 72 h at 125 rpm at 37° with 5% CO_2_ and 5% humidity.

Cells were then harvested at 4,000 *g* for 10 min and resuspended in 50 mL lysis buffer (50 mM HEPES pH 7.4, 1 M NaCl, 1 mM PMSF, 1 mM DTT, and 1 tablet of protease inhibitor (Sigma)). Lysis was performed using 15 loose and 15 tight plunges of a Wheaton glass dounce. The lysate was clarified using a 45 min, 360,000 *g* spin in a Ti70 rotor. The supernatant was incubated with 1 mL Streptactin sepharose beads (IBA Life Sciences) for 1 h. Beads were washed with 40 mL of lysis buffer followed by 30 mL labelling buffer (50 mM HEPES pH 7.4, 300 mM NaCl, 10 mM MgCl_2_, 1 mM EGTA, 10% glycerol, 1 mM DTT). Beads were then collected and incubated with purified SFP protein and an LD655 dye functionalized with CoA (Lumidyne) at room temperature for 30 min. Beads were washed with 30 mL labeling buffer. If proteins were to be kinase or phosphatase treated, they were additionally washed with 30 mL kinase (20 mM HEPES pH 7.5, 300mM NaCl, 10 mM MgCl2, 200 μM ATP, 10% glycerol, 1 mM DTT) or phosphatase (20 mM HEPES pH 7.5, 300mM NaCl, 1mM MnSO4, 10% glycerol, 1 mM DTT) buffer. Protein was eluted in 1 mL fractions in its final buffer supplemented with 10 mM desthiobiotin and concentrated using Amicon Ultra 30K concentrators. For kinase and phosphatase treatment, 5 μL of CKII kinase (New England Biolabs) and 2.5 μL λ phosphatase (New England Biolabs) was added per 50 μL concentrated protein, respectively, and the samples were incubated at 30 °C for 1 h. Final protein concentration was measured using Bradford reagent, and aliquots were snap-frozen in liquid nitrogen.

### In Vitro Transcription and RNA Labeling

For in vitro transcription of long viral RNA, the region of interest was first PCR amplified from a plasmid (N plasmid was a generous gift from the Krogan lab (*59*), 5’UTR plasmid was a generous gift from the Gladfelter lab (*52*), and SARS-CoV-2 cDNA plasmid was a generous gift of the Thiel lab (*51*)) using a forward primer with a T7 polymerase binding site (TAATACGACTCACTATAGGG). The amplified DNA was tested for purity on a 0.8% agarose gel and cleaned up using GlycoBlue and ethanol precipitation. RNA was generated using the HiScribe T7 Quick kit (New England Biolabs) and extracted using trizol and isopropanol precipitation. RNA was Cy3 labeled using a Label IT kit (Mirus Bio), and RNA purity was verified using a 0.8% agarose gel. All RNA structure predictions were done on the RNAfold server (*53, 54*).

### Sample Preparation and Microscopy

Purified protein and RNA samples were diluted into the imaging buffer (50 mM HEPES pH 7.4, 150 mM NaCl, 5 mM MgCl_2_, 1 mM EGTA, 1 mM DTT, 1% pluronic) to their final concentration and introduced into the flow chamber. Samples were settled onto the coverslip for 25 min before imaging. We confirmed that all of the condensates were settled to the surface within 25 min, as we did not observe an increase in the number of condensates per viewing area of the coverslip, and we could not detect freely diffusing condensates in the flow chamber after 25 min. Different buffers were used for phosphatase-treated (50 mM HEPES pH 7.4, 150 mM NaCl, 0.5 mM MnSO4, 1 mM DTT, 1% pluronic) and kinase treated (20 mM HEPES pH 7.5, 150 mM NaCl, 5 mM MgCl_2_, 100 μM ATP, 1 mM DTT, 1% pluronic) samples.

Imaging was performed using a custom-built fiber-coupled Nikon Ti-E Eclipse microscope equipped with an objective-type total internal reflection fluorescence (TIRF) illuminator and 100X 1.49 N.A. Plan Apo oil immersion objective (Nikon). The samples were excited in near-TIRF using 561 and 633 nm laser beams (Coherent). The fluorescent signal was detected by Andor Ixon electron-multiplying CCD (EMCCD) Camera (512×512 pixels). The effective pixel size was 160 nm after magnification. 10 single-frame images were acquired for each condition in each replicate.

### Image Analysis

To calculate the saturation concentration (c_sat_), the area of each condensate was quantified with Fiji using the Phansalkar function with a 30-pixel radius and a minimum condensate size of 10 pixels. The volume of the condensates was estimated from 2D projections by taking the semi-principal axis in the z-plane as the geometric average of semi-principal axes in the xy plane. The total volume of the condensates settled per micron squared on the coverslip was quantified. Conditions that resulted in measurable condensate volumes were fit to linear regression in Origin. The x-intercept of the linear regression represents c_sat_, the minimum protein concentration that results in condensate formation. The aspect ratio was calculated for individual condensates in Fiji using the Phansalkar function with a 30-pixel radius and a minimum condensate size of 10 pixels. To measure fusion times, condensates were visualized as they settled on the imaging surface from 5 to 25 min after mixing. Movies were recorded at 5 frames per second. Fusion times were calculated as the time between the last frame where two condensates appear separated (i.e., no overlap) and the first frame where the fused condensate appears nearly spherical (aspect ratio = 1.1).

### In Vitro FRAP

In vitro FRAP assays were performed with a Zeiss 880 Confocal Laser Scanning Microscope equipped with a 100x, 1.4 NA oil immersion objective. Samples were prepared as described above. All in vitro FRAP experiments used the 405, 488, 561, and 633 nm lasers at 100% for bleaching conditions. For N-polyC at 150 mM NaCl, 5 bleaching iterations were used, and images were taken every 2 s for 232 s. For N-SARS RNA at 150 mM NaCl, 100 bleaching iterations were used, and images were taken every 10 s for 8 min. For the experiments conducted at 50 mM NaCl, droplets containing polyC RNA or SARS RNA were bleached with 20 iterations, and images were taken every 5 s for 5 min. For droplets containing N protein only, 20 bleaching iterations were used, and images were collected every 2 s for 6.5 min. All data were acquired using Zeiss Zen 2.3 SP1 FP3 (black) (64bit) Version 14.0.20.201. Data were background corrected, converted to normalized intensity, and fit to an exponential decay function in Origin.

### Mass Spectrometry

The crosslinking analysis was performed as described by McGilvray et al. (*79*) with deviations outlined here. 15 μL 90 μM N protein in the labeling buffer was diluted 1:3 v/v into either water or labeling buffer. The diluted protein formed condensates in 100 mM NaCl in the final buffer, but not in the labeling buffer that contains 300 mM NaCl. Isotopically coded light (H12) and heavy (D12) BS3 (bis(sulfosuccinimidyl)suberate) crosslinkers (Creative Molecules Inc.) were immediately added to the diluted solution. Final concentration of BS3 was 0.8 mM for 100 mM NaCl dilution and 2.5 mM for 300 mM NaCl dilution. Crosslinking was performed for 30 min at room temperature and then quenched with 1 M Tris pH 8.0 buffer. This experiment was performed in duplicate, once with D12-BS3 crosslinking N protein in condensates and H12-BS3 crosslinking soluble N protein and once with the labels reversed. The crosslinked proteins from both channels were pooled before acetone precipitation. Protein was precipitated with acetone overnight at −20 °C. Protein was pelleted, the supernatant was removed, and the samples were air-dried for 10 min. The pellets were brought up in 8 M urea, reduced with TCEP, alkylated with iodoacetamide, diluted 4-fold, and digested with two rounds of trypsin. Peptides were desalted on a C18 MacroTrap column (Michrom Bioresources) and fractionated on a Superdex Peptide (GE Life Sciences) size-exclusion column. Fractions enriched in crosslinked peptides were dried and resuspended in 0.1% formic acid for mass spectrometry. Later eluting fractions were used for phosphorylation analysis. Each of the replicates was fractionated by size exclusion chromatography (SEC) and each fraction was injected twice, generating 16 mass spectrometry files for analysis.

Mass spectrometry was acquired on an Orbitrap Fusion Lumos coupled with an Easy-Spray nanoelectrospray ion source, a 15 cm × 75 μm PepMap C_18_ column (Thermo), and an M-class NanoAcuity UPLC system (Waters). Liquid chromatography mass spectrometry (LC-MS) runs were 90 min long. Precursor ions were measured in the Orbitrap at 120 k resolution. Selected precursor ions (triply charged and higher) were isolated, and the product ions were measured in the Orbitrap at 30 k resolution. Samples that were analyzed for phosphorylation were run similarly except only HCD product ions spectra were collected and doubly charge precursors were included.

Crosslinked spectra were identified with Protein Prospector 6.2.23 (*80*) using the combination of DSS/DSS:2H12 at uncleaved Lys residues and protein N-terminus as the crosslinking reagents. The corresponding light and heavy dead-end modifications, incorrect monoisotopic peak assignment (+1Da neutral loss), N-terminal pyroglutamate formation, methionine oxidation, and acetylation and loss of the protein N-terminal methionine were set as variable modifications with three variable modifications per peptide allowed. Trypsin with two missed cleavages was the digestion enzyme and the mass tolerances were 20 and 30 ppm for precursor and product ions. The search database comprised the 14 most abundant proteins (sorted by Spectral Abundance Factor) found in the linear peptide SEC fractions, alongside a decoy database that was ten times longer. Crosslink spectral matches were classified at a 1% false discovery rate (FDR) threshold.

Protein Prospector was used to search for phosphopeptides of the linear peptide fractions using the following parameters: tryptic specificity with 2 missed cleavages, 7 and 15 ppm precursor and product ion tolerances, and carbamidomethyl (C) fixed modification. Variable modifications were as above except that Phospho (STY) was included and only dead-end crosslink modifications were included. Peptides were reported with a maximum expectation value of 0.001, and a minimum prospector score of 15. A site localization in peptide (SLIP) score threshold of 6 was used to determine site-localization. All phosphopeptide spectra reported were manually inspected for evidence of correct site assignment. Where S or T residues are adjacent, the location of the phosphosite is ambiguous (see Table S2). All phosphorylation data can be accessed <u>here</u> (or on the MS-Viewer website with search key “jyovxenjny”).

A supplemental search of the crosslinked peak lists was made for crosslinks that also contained a phosphorylation site. This search was identical to the previous crosslinking searches except that Phospho (ST) was included as a variable modification, and the search was limited to the N protein sequence. Search results containing a phosphorylation site were manually assessed. Only phosphorylation sites that had been discovered in the phospho search of the linear peptides were considered.

For quantitating the isotopically labeled crosslinks, peak areas were measured from the extracted precursor ion chromatograms (XICs) using the small molecule interface of Skyline (v20.1.0.155). A Skyline transition list was generated containing the elemental composition of each distinct peptide pair with both light and heavy BS3 modification and in each charge state detected in the Prospector search. Retention times were present in the transition list to help with peak detection. Peptide level measurements were imported into R and summarized at the level of unique residue pairs (“crosslinks”). For each precursor ion, the log_2_ ratios of heavy to light peptides were calculated and mean normalized to 0 for each of the two biological replicates and then transformed into log_2_ (water/salt) ratios. Weighted t-tests were performed in R.

### Drug Screening and Image Processing

For the FDA-approved drug screen, 75 mL of N protein at 16 μM was purified from HEK293S GNTI-cells. polyC RNA was obtained from Sigma. The FDA-Approved Drug Library (TargetMol) has 1,200 compounds of well-characterized biological activity. 10 mM compounds were stored in 100% dimethyl sulfoxide (DMSO) in 384 well plates. For screening plates, 20 μL imaging buffer (50 mM HEPES pH 7.4, 150 mM NaCl, 5 mM MgCl_2_, 1 mM EGTA, 1 mM DTT, 1% pluronic) was aliquoted into 384 well, glass-bottomed plates (Greiner Bio-One) and incubated for 5 min. Buffer was removed, and 0.5 μL of 2 mM compounds were stamped into 384 well plates with an Analytik-Jena Cybio Well Vario liquid handler. The final concentration of 7.8 μM N protein and 50 ng/μL polyC RNA were added to each plate. Each well contained a 25 μL mixture and 40 μM compound in the primary screen.

Wells were homogenized with a Bioshake 3000 ELM orbital shaker at 2,400 rpm for 45 s and condensates were allowed to settle for 1 h before imaging. 4 images (each 224 × 167 μm) were taken at 40x with a Molecular Devices ImageXpress Micro High Content Imaging System. Images were analyzed with Metamorph Imaging software calculating condensate count and area. The volume of the condensates was calculated as described above. Data was then uploaded to CCDvault for normalization to DMSO vehicle control wells. Wells that exhibit 3 standard deviations from the untreated sample and as well as others that produced qualitative morphological changes in N condensates were selected for dose-response screening. The same N protein and RNA were tested against candidate drugs using a 10-point serial dilution starting at 40 μM using the same procedure. Dose-response curves were generated by fitting to the equation:

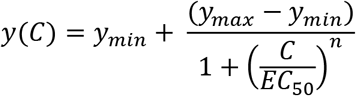

where *C* is the concentration of the drug, *n* is the Hill coefficient, *EC_50_* is the half-maximal response concentration, and *y* is either the volume of the N condensates or the number of N condensates settled onto per micron squared area on the coverslip.

### Dose-Response with SARS-CoV-2 Infected Cells

Vero-E6 (ATCC, CRL-1586) and Calu-3 cells (ATCC HTB-55) were cultured in high glucose DMEM (Gibco) supplemented with 10% FBS (R&D Systems), 1X GlutaMAX (Gibco), and 1X PenStrep (Gibco) at 37°C and 5% CO_2_. For screening selected drugs against infected cells, 2500 Vero-E6 (12 μl/well) or 10000 Calu-3 (12 μl/well) were seeded in 384-well white optical-bottom tissue culture plates (Nunc) with the Multidrop Combi liquid handling instrument (Thermo Fisher Scientific). Cells were allowed to recover for 24 h for Vero-E6 and 48 h for Calu-3 at 37°C and 5% CO_2_. Dose responses were generated by diluting the compounds using a Cybio Well Vario liquid handler (Analytik Jena), leading to a final concentration of DMSO at 0.4% in the assay plate (v/v). Cells were incubated at 37°C and 5% CO_2_ for 1 h before infection. The viral inoculum was prepared such that the final MOI = 0.05 upon addition of 6 μl/well viral inoculum. After complete CPE was observed in DMSO-treated, infected wells (72 h post-infection (hpi) for Vero-E6 and 96 hpi for Calu 3), the plates were developed with the CellTiter-Glo 2.0 reagent (Promega) according to the manufacturer’s instructions. For Vero-E6, the reagent was diluted 1:1 (v/v) in PBS. Luminescence of developed plates was read on a Spectramax L (Molecular Devices). Each plate contained 24 wells uninfected/DMSO treated cells (100% CPE inhibition) and 24 wells infected/DMSO treated cells (0% CPE inhibition). Average values from those wells were used to normalize data and determine % CPE inhibition for each compound well. To determine the cytotoxicity of the compounds, the same protocol was used but with 6 μL growth media added instead of viral inoculum. The data were plotted and analyzed in Origin and fit to the dose-response function above.

### HEK293T Cell Culture and Transfection

HEK293T (RRID: CVCL_0063) cells were cultured in phenol-negative DMEM media supplemented with 10% FBS and 1% PS at 37° with 5% CO_2_. Prior to imaging, cells were transferred to glass-bottomed plates (Nunc Lab-Tek, 0.4 mL working volume) at ~25% confluence and allowed to recover for 24 hours. Media was exchanged into phenol-negative DMEM media supplemented with 10% FBS, and cells were transfected with N-GFP, NΔR2-GFP, or GFP only constructs. For each well, 400 ng of DNA was added to 20 μL DMEM media. 1.2 uL FuGENE HD transfection reagent (Promega) was added, and the mixture was incubated at room temperature for 15 minutes before being added to the cell culture. Cells were allowed to express the constructs for 4 days. 1 hour prior to imaging, DRAQ5 stain at 1:1000 was added to the media.

### In Vivo Imaging and FRAP Assays

In vivo imaging and FRAP measurements were performed with a Zeiss 880 Confocal Laser Scanning Microscope equipped with a 63x, 1.4 NA oil immersion objective. Samples were prepared as described above. All in vivo FRAP experiments used the 405, 488, 561, and 633 nm lasers at 100% for bleaching conditions. Cells expressing N-GFP, ΔR2-GFP, or GFP-only were bleached with 30 iterations, and images were collected continuously at 6.7 Hz for 30 s. All data were acquired using Zeiss Zen 2.3 SP1 FP3. Data were converted to the normalized intensity and fit to an exponential decay function in Origin.

### Data Availability and Transparency

Raw data and software used in this study are uploaded to Github.

## Author Contributions

A.J., L.F., M.J.T., E.W., J.S., and A.Y. conceived of experiments. A.J. cloned the constructs and generated RNA. A.J. and L.F. purified the protein and performed the in vitro experiments. M.J.T. performed the mass spectrometry experiments. A.J. cultured the cells and performed live-cell imaging for the N expression experiments. X.N. cultured the cells for the SARS-CoV-2 infection dose-response curves. A.J., E.W., and X.N. performed drug screening experiments. A.J., M.J.T., E.W., and A.N. analyzed the data. K.C. created the model figures. A.J., L.F., M.J.T., E.W., A.N., and A.Y. wrote the manuscript with inputs from all authors.

## Acknowledgments

We thank John T. Canty and other members of Yildiz laboratory for helpful discussions, the UC Berkeley Biological Imaging Facility for use of their confocal FRAP microscope, Denise Schichnes for FRAP advice and training, Thomas Graham (UC Berkeley) for advice and reagents for in vitro transcription, Kathy Li (UCSF) for assisting with CLMS sample preparation, Nevan Krogan (UCSF), Amy Gladfelter (UNC) and Volker Thiel (Univ. Bern) for providing the plasmids, the UC Berkeley MacroLab for TEV protease and competent cells, and the UC Berkeley Cell Culture Facility for HEK293S GNTI- and HEK293T.

